# Accurate Bayesian inference of sex chromosome karyotypes and sex-linked scaffolds from low-depth sequencing data

**DOI:** 10.1101/2023.09.15.557900

**Authors:** Madleina Caduff, Raphael Eckel, Christoph Leuenberger, Daniel Wegmann

## Abstract

The identification of sex-linked scaffolds and the genetic sex of individuals, i.e. their sex karyotype, is a fundamental step in population genomic studies. If sex-linked scaffolds are known, single individuals may be sexed based on read counts of next-generation sequencing data. If both sex-linked scaffolds as well as sex karyotypes are unknown, as is often the case for non-model organisms, they have to be jointly inferred. For both cases, current methods rely on arbitrary thresholds, which limits their power for low-depth data. In addition, most current methods are limited to euploid sex karyotypes (XX and XY). Here we develop `BeXY`, a fully Bayesian method to jointly infer the posterior probabilities for each scaffold to be autosomal, X-or Y-linked and for each individual to be any of the sex karyotypes XX, XY, X0, XXX, XXY, XYY and XXYY. If the sex-linked scaffolds are known, it also identifies autosomal trisomies and estimates the sex karyotype posterior probabilities for single individuals. As we show with downsampling experiments, `BeXY` has higher power than all existing methods. It accurately infers the sex karyotype of ancient human samples with as few as 20,000 reads and accurately infers sex-linked scaffolds from data sets of just a handful of samples or with highly imbalanced sex ratios, also in the case of low-quality reference assemblies. We illustrate the power of `BeXY` by applying it to both whole-genome shotgun and target enrichment sequencing data of ancient and modern humans, as well as several non-model organisms.

## 2 Introduction

Many population genomic analyses rely on an accurate identification of the genetic sex of the individuals as well as an identification of sex-linked scaffolds. The identification of genetic sex is used to reveal patterns of sex-specific ecology and evolution, for example concerning behaviour and social structure (e.g. Pečnerová et al., 2017; Gower et al., 2019), foraging ecology (e.g. Louis et al., 2021) or sex-biased dispersal and philopatry (e.g. Bidon et al., 2014; Herrero et al., 2021). In the context of ancient humans cultures, the identification of the genetic sex has been used to study social structure (e.g. Žegarac et al., 2021; Hedenstierna-Jonson et al., 2017), religion (e.g. De La Cruz et al., 2008), sex-biased migration (e.g. Veeramah et al., 2018) or to evidence women participating in prehistorical battles (e.g. Burger et al., 2020). The identification of sex-linked scaffolds further allows to study sex-linked disease susceptibility (Schurz et al., 2019), sex-biased gene expression (Grath and Parsch, 2016) or sex-linked gene-flow (Bidon et al., 2014). In addition, sex-linked scaffolds often require special treatment in population genomic analyses, for example for fair filtering on read depth and minor allele frequency, genotype calling and Hardy-Weinberg checks, but also because they bias estimates of genetic diversity such as heterozygosity (Ellegren, 2009), estimates of mutation rates (Ellegren, 2007), genome-wide association studies (Gao et al., 2015) and assessments of population genetic structure (Benestan et al., 2017). If genetic sex and/or sex-linked scaffolds are unknown, they need to be characterized using dedicated methods that can be easily integrated in population genomic analyses.

The mechanisms of sex determination are remarkably variable (The Tree of Sex Consortium et al., 2014), and genetic sex determination through sex chromosomes has evolved independently many times throughout eukaryotes (Graves, 2008). However, two major systems can be outlined. In therian mammals, beetles, many flies and some fish, males are heterogametic (XY) and females are homogametic (XX). In contrast, in birds, snakes, butterflies and some other fish, females are heterogametic (ZW) and males are homogametic (ZZ) (Graves, 2008; Bachtrog et al., 2014; Stöck et al., 2021). Visual identification of the sex based on morphology or behaviour can be challenging or even impossible (Fairbairn et al., 2007), for example for species without sexual dimorphism (Kocijan et al., 2011), but also for juveniles (Kocijan et al., 2011), ancient or historical samples (Buonasera et al., 2020), forensic samples (e.g. hair, Madel et al., 2016), as well as samples of unknown origin, e.g. environmental samples or faeces (Peppin et al., 2010; Sastre et al., 2009).

Molecular sexing through genetic data is an alternative to sexing from observations as it is accurate, simple, cheap, time-efficient, applicable to any tissue and can often piggyback on other genetic analyses (Buonasera et al., 2020). For example, molecular sexing from next-generation sequencing data exploits the difference in sex chromosome ploidy between males and females: in XY-systems, roughly half as many reads are expected to map to the X-chromosome in males than females, and no reads are expected to map to the Y-chromosome in females (Palmer et al., 2019). The same logic applies to ZW-systems. These unique mapping patterns can be used i) to deduce the genetic sex of individuals (sexing), ii) to deduce the type (i.e. autosomal or sex-linked) of scaffolds or iii) to deduce both jointly in case both are unknown. In the following, we describe these three major scenarios.

Current methods to sex individuals assume a reference genome of which the scaffold types are known. A typical case is the sexing of ancient human samples, for which two main methods are currently used: *R*_*y*_ and *R*_*x*_. *R*_*y*_ (Skoglund et al., 2013) calculates the fraction of reads mapping to the Y-chromosome relative to the total number of reads mapping to both sex chromosomes (X and Y). *R*_*x*_ (Mittnik et al., 2016) calculates the average of the fraction of reads mapping to the X chromosome relative to the number of reads mapping to each autosome. Both methods then assign the sex based on thresholds determined on small data sets (30 and 20 samples, respectively). Nonetheless, most current studies (e.g. Allentoft et al., 2015; Scheib et al., 2018; Margaryan et al., 2020; Furtwängler et al., 2020; Lipson et al., 2022; Liu et al., 2022b; Reitsema et al., 2022) use the same thresholds for sexing without considering differences in sequencing methods, sequencing depth or sample treatment.

The methods *R*_*y*_ and *R*_*x*_ are also both limited to the euploid karyotypes XY and XX, although sex karyotypes aneuploidies in humans are relatively common with an estimated incidence of 1 in 440 births (Breman and Stankiewicz, 2021). Several studies that used large cohorts detected aneuploid individuals, but only by manually inspecting individuals with ambiguous mapping statistics (Rivollat et al., 2020; Villalba-Mouco et al., 2021; Ebenesersdóttir et al., 2018). To our knowledge, there exists currently only a single method that can identify sex aneuploidy in humans (`seGMM`, Liu et al., 2022a), and it does so based on the mean and standard deviation of normalized mapping counts to X and Y using predefined cutoffs.

In the second scenario, the goal is to infer the scaffold types, assuming that the genetic sex of the individuals is known. Scaffold types are often unknown for non-model organisms with low-quality and fragmented draft genome assemblies due to high cost and challenges associated with complete genome assembly (Ellegren, 2014). Approaches to identify sex-linked scaffolds include synteny-based alignments to closely related species with a chromosomal-level genome assembly (Grabherr et al., 2010) or comparison of the sequencing depth of each scaffold between males and females (Palmer et al., 2019). The tool `findZX` (Sigeman et al., 2022) implements a pipeline to identify X*/*Z chromosomes through differences in sequencing depth and heterozygosity. The method `discoverY` identifies Y-linked scaffolds through differences in sequence similarity and sequencing depth between males and females (Rangavittal et al., 2019).

In the third scenario, both the sex karyotypes of individuals as well as the scaffold types are unknown, as is often case for non-model organisms. The method `SATC` (Nursyifa et al., 2022) infers both sex and sex-linked scaffold in a two step procedure. It first conducts a principal component analysis (PCA) on normalized depth and uses Gaussian mixture clustering to identify males and females. It then uses a *t*-test to identify sex-linked scaffolds as those for which the normalized depth differs between the two sex groups. However, `SATC` relies on hard thresholds and can not identify Y-linked scaffolds. The method `SeXY` (Cabrera et al., 2022) uses a synteny-based approach to identify sex-linked scaffolds and performs sexing based on the X to autosome depth ratios. While `SeXY` can perform sexing with a single individual at low depth and was tailored to also work with reference genomes of divergent taxa, its sexing is based on arbitrary thresholds. Other methods (Robledo-Ruiz et al., 2023; Gautier, 2014) can also infer both sex and sex-linked loci but are designed for SNP data sets (e.g. RAD sequencing) and classify each locus individually. None of the tools allows for aneuploid sex karyotypes.

Here, we present a Bayesian method designed to work in all three scenarios described above. Our method, which we call *Bayesian estimation of X and Y*, or `BeXY` for short, jointly infers the type of all scaffolds as well as the sex karyotype for all individuals. In contrast to existing methods, `BeXY` i) does not rely on hard thresholds but instead infers probabilities for scaffold types and sex karyotypes, ii) is flexible in respect of incorporation of existing knowledge on sex-linked scaffolds and sex karyotypes in the model, iii) can identify both X- and Y-linked scaffolds, iv) can detect aneuploid sex karyotypes, v) can detect trisomies for scaffolds known to be autosomal and vi) retains high accuracy for low-depth and highly fragmented genome assemblies. To illustrate our method, we applied it to whole-genome shotgun and target enrichment sequencing data of ancient and modern humans as well as to data from several non-model organisms. Using simulations, we show that our method outperforms current methods in all scenarios.

## 3 Materials and Methods

### 3.1 The model

For readability, we will present the model using the XY notation, although the model applies equally well to ZW-systems.

Consider a reference genome consisting of *C* scaffolds. Let *n*_*rc*_ denote the number of sequencing reads of sequencing run *r* = 1, …, *R* mapped to scaffold *c*. These mapping statistics can be obtained easily from indexed BAM-files either with the `idxstats` command in `Samtools` (Li et al., 2009) or with the `BAMDiagnostics` command in `ATLAS` (Link et al., 2017). Each sequencing run *r* = (*i, m*) corresponds to a certain individual *i* = 1, …, *I* sequenced with a specific method *m* = 1, …, *M* such as, for instance, target enrichment sequencing, whole-genome shotgun sequencing or restriction site associated DNA sequencing (RAD). These methods likely differ in their mapping characteristics and are thus modeled with their own set of parameters. Individuals may have been sequenced with multiple runs and potentially different sequencing methods and the model described here allows to make use of all that data jointly.

We model the vector of counts ***n***_*r*_ = (*n*_*r*1_, …, *n*_*rC*_) of sequencing run *r* using a Dirichlet-Multinomial (DM) distribution (Johnson et al., 1997) as

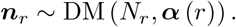

Here, *N*_*r*_ = ∑_*c*_ *n*_*rc*_ is the total number of mapped reads of run *r* and the parameter vector ***α*** (*r*) = (*α*_1_ (*r*), …, *α*_*C*_ (*r*)) models both the expected number and the variance of reads mapping to each scaffold.

We model *n*_*rc*_ of run *r* = (*i, m*) using two major components: the ploidy *p*_*ic*_ of scaffold *c* in individual *i*, and some method- and scaffold-specific mapping attractor *γ*_*mc*_. These mapping attractors may differ among scaffolds due to, for instance, scaffold length or mappability as more reads are expected to map to longer scaffolds or to scaffolds with fewer repetitive regions. We model *α*_*c*_(*r*) as

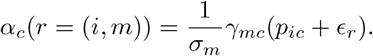

Here, *ϵ*_*r*_ is an run-specific parameter that allows for noise due to erroneously mapped reads and *σ*_*m*_ is a positive scaling parameter that models the variance of the Dirichlet-Multinomial distribution. To avoid non-identifiability issues with *σ*_*m*_, we impose the constraint that 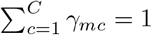.

#### Ploidy

The ploidy *p*_*ic*_ depends on two factors: the sex karyotype *s*_*i*_ of individual *i* and the scaffold type, i.e. whether a scaffold is autosomal, X or Y-linked. We distinguish seven sex karyotypes, which correspond to the most frequent sex karyotypes observed in humans: the heterogametic sex (*s*_*i*_ = XY), the homogametic sex (*s*_*i*_ = XX), the monosomy *s*_*i*_ = X0 as well as the three trisomies *s*_*i*_ = XXY, *s*_*i*_ = XYY and *s*_*i*_ = XXX and the tetrasomy *s*_*i*_ = XXYY. We will denote by 𝒜 = {X0, XXY, XYY, XXX, XXYY} the set of considered aneuploid sex karyotypes.

In theory, the ploidy is given by the number of copies of each scaffold, for example *p*_*ic*_ = 2 for all autosomal scaffolds or *p*_*ic*_ = 1 for both X and Y-linked scaffolds of an individual with sex karyotype XY. However, in low-quality reference genomes, the ploidy ratio between the sex karyotypes often differs from the canonical cases due to noise (Nursyifa et al., 2022). We here model the ploidy using a continuous, scaffold-specific parameter *ρ*_*c*_ ∈ [0, 1] such that *ρ*_*c*_ = 0 corresponds to a canonical Y, 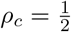 to a canonical autosome, and *ρ*_*c*_ = 1 to a canonical X. Intermediate values of *ρ*_*c*_ then signify ploidies that are different from these cases, for instance for a misassembled scaffold that consists of both autosomal and gonosomal sequences. We model the resulting ploidies *p*_*ic*_ for all considered sex karyotypes as a function of *ρ*_*c*_ as linear functions between the canonical cases as given in Table 1 and illustrated in Figure 1.

**Table 1:**
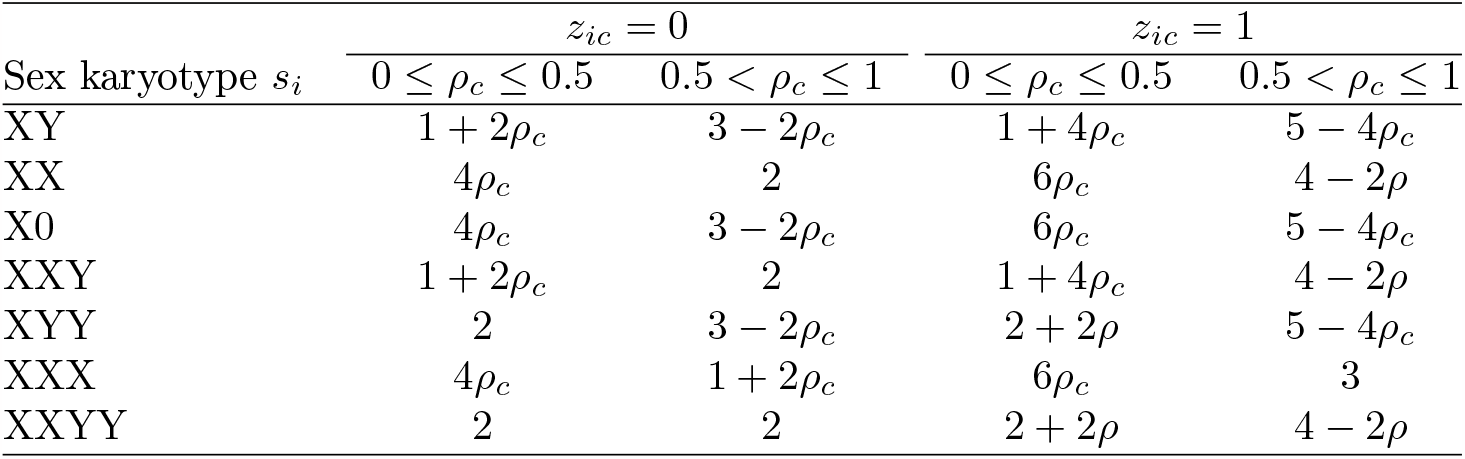
Ploidies *p*_*ic*_ as a function of ploidy parameter *ρ*_*c*_ for all sex karyotypes. The first two columns with *z*_*ic*_ = 0 are for the autosomal euploid case, while the latter columns with *z*_*ic*_ = 1 are for the autosomal triploid case.

**Figure 1.**
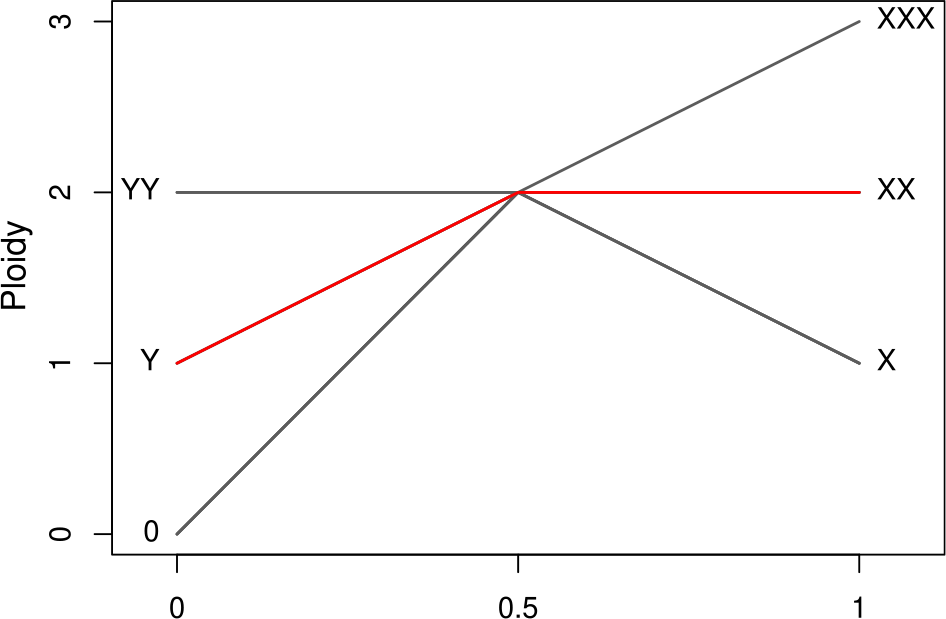
Ploidy as a function of *ρ*_*c*_ for all sex karyotypes, obtained by following the line from the number of Y copies to the number of X copies of that karyotype. The red line exemplifies the ploidy function for the karyotype XXY.

`BeXY` classifies each scaffold into one of four categories: autosomal (*t*_*c*_ = A), X-linked (*t*_*c*_ = X), Y-linked (*t*_*c*_ = Y) or “different” (*t*_*c*_ = D) for ploidies that differ from the canonical cases. We achieve this by modeling *ρ*_*c*_ through a mixture of four Beta distributions. For autosomes, we use a symmetrical Beta distribution *ρ*_*c*_|*t*_*c*_ = A ∼ Beta(*µ, µ*), which has an expected value 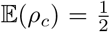. For X-linked scaffolds, we use a left-skewed distribution *ρ*_*c*_|*t*_*c*_ = X ∼ Beta(*α*, 1) with *α <* 1, which has an expected value close to one. For Y-linked scaffolds, analogously, we use a right-skewed distribution *ρ*_*c*_|*t*_*c*_ = Y ∼ Beta(1, *β*) with *β <* 1, which has an expected value close to zero. Finally, for non-canonical scaffolds of type *t*_*c*_ = D we use an uninformative Beta distribution *ρ*_*c*_|*t*_*c*_ = D ∼ Beta(1, 1) (see the Supplementary Materials for a more detailed description of the parametrization).

#### Distribution of karyotypes

We assume that the distribution of sex karyotypes *s*_*i*_ in a sample are described well by the frequency *a* of aneuploids and the frequency *f* of XX to XY karyotypes among euploids, such that

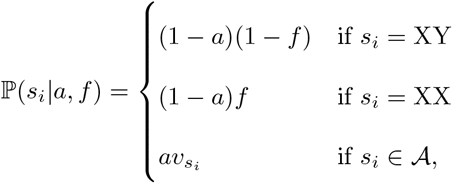

where ***v*** = (*v*_X0_, …, *v*_XXYY_) correspond to the relative frequencies of aneuploid karyotypes, with ∑_*s*∈*𝒜*_ *v*_*s*_ = 1. By default, we set ***v*** to the relative frequencies observed in humans (Skuse et al., 2018).

#### Autosomal trisomies

It is straightforward to extend the `BeXY` model to account for autosomal trisomies. We introduce a binary parameter *z*_*ic*_ that indicates whether individual *i* at scaffold *c* is diploid (*z*_*ic*_ = 0) or triploid (*z*_*ic*_ = 1). The ploidies *p*_*ic*_ for both cases are given in Table 1, where *z*_*ic*_ = 0 results in a ploidy *p*_*ic*_ = 2 and *z*_*ic*_ = 1 results in a ploidy *p*_*ic*_ = 3 at the canonical autosomes with 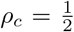. To avoid the need for model choice, we allow *z*_*ic*_ = 1 only for scaffolds known to be autosomal (i.e. with 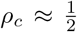). Since autosomal trisomies are lethal for most scaffolds, the user further needs to specify for which scaffolds trisomies are to be considered.

We assume *z*_*ic*_ ∼ Bernouilli(*π*) and set 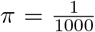 by default, which matches the frequency of trisomy 21 in humans.

### 3.2 Bayesian inference

We use a Markov chain Monte Carlo (MCMC) scheme to generate samples from the posterior ℙ(***s, ρ, γ, ϵ, σ, z***|***N***), where ***N*** denotes the *R × C* matrix of read counts with entries [***N***]_*rc*_ = *n*_*rc*_. We update *z*_*ic*_ and *t*_*c*_ using Gibbs sampling and all other parameter using standard Metropolis-Hastings updates. See Supplementary Figure 1 for a Directed Acyclic Graph (DAG) illustrating the full model and the Supplementary Material for a description of all prior distributions and MCMC updates used.

### 3.3 Implementation

`BeXY` is a command-line tool implemented in `C++` making use of the MCMC framework of the statistical library `stattools` (bitbucket.org/wegmannlab/stattools). It implements three tasks: `infer`, `inferScaffolds` and `sex`. The task `infer` is used to infer sex karyotypes and scaffold types jointly, the task `inferScaffolds` is used to infer scaffold types if sex karyotypes are known, and the task `sex` is used to infer sex karyotypes, i.e. only the individual- and run-specific parameters *s*_*i*_ and *ϵ*_*r*_, while using estimates of ***ρ, γ***_*m*_ and *σ*_*m*_ previously learned from a larger data set. `BeXY` reads mapping statistics obtained with either the `idxstats` command in `Samtools` (Li et al., 2009) or with the `BAMDiagnostics` command in `ATLAS` (Link et al., 2017). It is open-source and freely available through the `git`-repository at bitbucket.org/wegmannlab/bexy, along with a wiki detailing its usage and a custom R-package called `bexy` to visualizes the results of `BeXY` and to classify individuals based on custom thresholds on the posterior probabilities of the sex karyotypes.

### 3.4 Application to empirical data sets

#### Ancient humans

We downloaded publicly available data of ancient humans sequenced with two different methods. We first downloaded 954 individuals sequenced with Illumina whole-genome shot-gun (WGS) sequencing (Antonio et al., 2019; Damgaard et al., 2018; Ebenesersdóttir et al., 2018; Lazaridis et al., 2014; Scheib et al., 2018; Hofmanová et al., 2016; Broushaki et al., 2016; Gamba et al., 2014; Günther et al., 2018; Margaryan et al., 2020; De Barros Damgaard et al., 2018; Marchi et al., 2022; Jones et al., 2017, 2015) in fastq-format and processed these with the `Gaia` part of the `ATLAS`-pipeline (bitbucket.org/wegmannlab/atlas-pipeline). Specifically, we trimmed the reads using `Trim Galore` (github.com/FelixKrueger/TrimGalore) with no quality filter and a length filter of 30. Reads were subsequently aligned to the GRCh38 human reference genome (NCBI RefSeq assembly GCF_000001405.26) using `BWA-MEM` (Li, 2013). Reads with a mapping quality below 30 were filtered out with `Samtools` (Li et al., 2009). Unmapped and, in the case of paired-end sequencing data, unpaired reads were removed with `Samtools`. Samples consisting of multiple libraries were merged using `Samtools`. Duplicate reads were marked using `picard-tools` MarkDuplicates (broadinstitute.github.io/picard). We further downloaded 2, 457 individuals sequenced with 1240k target enrichment capture sequencing (Fernandes et al., 2020; Fowler et al., 2022; Fu et al., 2016; Harney et al., 2021, 2022; Kennett et al., 2022; Lazaridis et al., 2022, 2016, 2017; Lipson et al., 2017, 2022; Liu et al., 2022b; Mathieson et al., 2015, 2018; Narasimhan et al., 2019; Novak et al., 2021; Olalde et al., 2018, 2019; Patterson et al., 2022; Prendergast et al., 2019; Reitsema et al., 2022; Rivollat et al., 2020; Sirak et al., 2021; Tiesler et al., 2022; Villalba-Mouco et al., 2021) directly in BAM-format. For both data sets, we used the task `BAMDiagnostics` in `ATLAS` (Link et al., 2017) to count the number of reads that aligned to each chromosome, ignoring duplicates and reads with a mapping quality below 30. The resulting files were used as an input for `BeXY`. For each data set, we ran `BeXY` with the task `infer` and setting `--allowTrisomyForAutosomes 21` to identify cases of trisomy 21. To investigate the effect of the prior choice on aneuploid sex karyotypes, we additionally ran `BeXY` with the task `sex`, using the inferred parameters ***ρ, γ***_*m*_ and *σ*_*m*_ and different values of 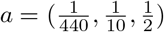.

#### Modern humans

We downloaded publicly available CRAM-files of 276 modern humans from the Simons Genome Diversity Project (SGDP, Mallick et al., 2016). Of these, 260 samples were sequenced using a PCR-free library preparation and 16 samples were sequenced using a PCR-based library preparation (Mallick et al., 2016). We used the `idxstats` command in `Samtools` to generate the input needed for `BeXY`.

#### Non-model organisms

We ran `BeXY` on the six WGS data sets of varying assembly quality provided by Nursyifa et al. (2022): impala (*Aepyceros melampus*), muskox (*Ovibos moschatus*), waterbuck (*Kobus ellipsiprymnus*), grey whale (*Eschrichtius robustus*), leopard (*Panthera pardus*) and the Darwin’s finches species complex encompassing 15 species. We used the `idxstats` files provided by (Nursyifa et al., 2022) and ran `BeXY` and `SATC` on these data sets. We used the default filters of `SATC` for both `BeXY` and `SATC`, i.e. we removed scaffolds with *<* 100kb length and a normalized depth outside the range (0.3, 2).

### 3.5 Downsampling experiments

The performance of `BeXY` likely depends on the sequencing depth of the individuals and, in case scaffold types are inferred, also on the sample size, the sex ratio among the samples, and the quality of the reference genome assembly. We tested the limits of `BeXY` and competing methods with respect to each of these challenging factors using dedicated downsampling experiments based on a subset of the ancient WGS data set consisting of all individuals with a depth *>* 2*×* (*n* = 116) that were also all identified as euploid on the full data sets.

#### Sexing individuals: euploids

To evaluate the robustness of sexing individuals with respect to low-depth data of euploid individuals, we downsampled (with replacement) the read counts of all individuals in the subset to various depth ranging from 200,000 to 100 reads per individual. We ran `BeXY` with the task `sex` on 100 replicates of downsampled counts, and set the prior parameters on *s* to the values expected for a human data set, 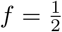 and 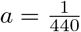 (Breman and Stankiewicz, 2021). We used parameters ***ρ, γ***_*m*_ and *σ*_*m*_ inferred from a single run on the full data of all *n* = 116 individuals and considered all classifications with state posterior probabilities P(*s*_*i*_|***n***_*r*_) *>* 0.9 as confident. We then compared the sex inferred from the downsampled data set with that inferred from the full data set. We also ran the two competing sexing methods *R*_*x*_ (Mittnik et al., 2016) and *R*_*y*_ (Skoglund et al., 2013) on the same data set using default parameters and counted the classifications “Sex assignment: The sample could not be assigned” (for *R*_*x*_) as well as “Not Assigned” (*R*_*y*_) as uncertain and all others as confident.

#### Sexing individuals: aneuploids

To investigate the power of the task `sex` to detect aneuploid individuals at various depths, we simulated aneuploid individuals based on the samples in the subset. We used individuals with karyotype XY to simulate samples with karyotypes XXY, XYY and XXYY and individuals with karyotype XX to simulate samples with karyotypes X0 and XXX as follows: Let *f*_*x*_ and *f*_*y*_ denote the relative change in ploidies between the karyotype to simulate and the karyotype of the individual serving as template (e.g. *f*_*X*_ = 2, *f*_*Y*_ = 2 to simulate the karyotype XXYY from an XY individual, or *f*_*X*_ = 0.5, *f*_*Y*_ = 1 to simulate a karyotype X from an XX individual). We then sampled reads (with replacement) from the vector ***n***_*r*_ according to the probabilities

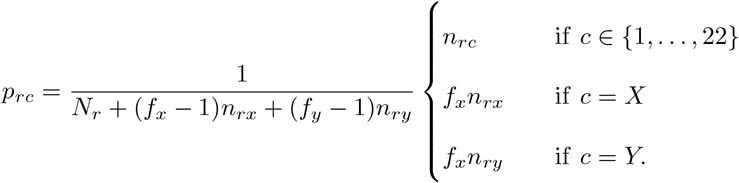

Using this procedure, we simulated samples for various depth ranging from 200,000 down to 100 reads per sample. Using the same parameters ***ρ, γ***_*m*_ and *σ*_*m*_ as for euploids, we ran `BeXY` with the task `sex` on 100 replicates of downsampled counts and identified confident classifications as described above, both on a per-karyotype basis as well as for euploid and aneuploid karyotype classes, ensuring that all karyotype were equally frequent within a class. Since the prior on the number of aneuploid samples, *a*, has a large impact on the classification accuracy of euploid and aneuploid individuals, we ran `BeXY` with both 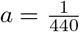 and 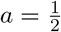.

We ran the competing sexing method `seGMM` (Liu et al., 2022a) on the same data set. Since the available implementation of `seGMM` requires a VCF file to perform the clustering of the samples for subsequent sexing, we re-implemented their approach and calculated the relevant statistics needed for sexing (the mean and standard deviation of normalized read counts for the X and Y chromosome for XX and XY samples) based on the full counts. While re-implementing, we also fixed an obvious bug in their code that led to a misclassification of XXX individuals (XXX karyotypes are expected to have 1.5 times the number of reads mapping to the X-chromosome than XX karyotypes, not twice as many). We counted all classifications that could not be assigned as uncertain and all others as confident.

#### Identifying autosomal trisomies

To investigate the power to detect autosomal trisomies at various depths, we simulated individuals with trisomy 21 based on the samples in the subset. We sampled reads (with replacement) from the vector ***n***_*r*_ as describes above and according to the probabilities

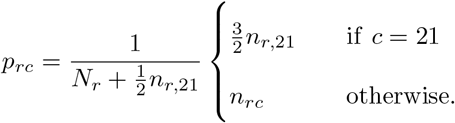

Using this procedure, we simulated samples with trisomy 21 for various depth ranging from 200,000 down to 100 reads per sample. We ran `BeXY` with the task `sex` on 100 replicates of these downsampled counts, setting `--allowTrisomyForAutosomes 21` to allow for trisomies on chromosome 21 and using the same parameters ***ρ, γ***_*m*_ and *σ*_*m*_ used above to sex individuals. To test for false positive classifications, we additionally downsampled counts without simulating any trisomies as described above and analysed them with `BeXY`. We then considered all classifications with state posterior probabilities P(*z*_*ic*_|***n***_*r*_) *>* 0.9 as confident. Since the prior on the number of samples with autosomal trisomy, *π*, has a large impact on the classification accuracy, we ran `BeXY` with both 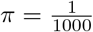 and 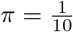.

#### Inferring scaffold types

We also investigated the power to infer sex karyotypes scaffold types jointly, using the same downsampled counts as for the first downsampling experiment described above (only euploids). We ran `BeXY` with the task `infer` and identified confident classifications as described above. We ran the competing method `SATC` (Nursyifa et al., 2022) on the same data set. For comparability with `BeXY`, we turned off the normalized depth range filter by setting it to (0, 100) in `SATC`. `SATC` does not provide an estimate of classification uncertainty and we assumed all classifications as confident. But we note that `SATC` throws an error if no good candidates for sex scaffolds were found through clustering or when one of the inferred sex groups only had one member. We counted these cases as uncertain.

To evaluate the robustness of our method to small data sets or those with large sex ratio imbalances when jointly inferring sex karyotypes and scaffold types, we first downsampled all individuals in the subset to 20K reads and then randomly compiled data sets with various XY:XX proportions differing in size and imbalance. We ran `BeXY` and `SATC` as described above.

To evaluate the robustness of our method with respect to low-quality reference genome assemblies, we simulated a low-quality reference genome based on the human reference genome GRCh38. To match the roughly exponential distribution of empirical low-quality reference genomes (e.g. waterbuck from Nursyifa et al., 2022), we defined length categories of 10^6^, 5 · 10^5^, 2 · 10^5^, 10^5^, 5 · 10^4^, 2 · 10^4^ and 10^4^ and split the chromosomes into scaffolds of these lengths. To limit mapping issues, we excluded the telomeres (15kb at each end of each chromosome), the pseudo-autosomal regions (PAR) on the X and Y chromosome as well as the centromeres of all chromosomes (all downloaded from ncbi.nlm.nih.gov/grc/human), with an additional buffer of 100kb around each of these regions. After splitting, we further excluded all resulting scaffolds that overlapped by more than 90% with gap regions consisting of N (hgdown-load.soe.ucsc.edu/goldenPath/hg38/database/gap.txt.gz, Nassar et al., 2023) as well as all scaffolds that overlapped by more than 90% with regions listed in the ENCODE Blacklist (Amemiya et al., 2019). We then re-mapped the sequencing data of all samples against this artificially created low-quality reference genome. To reduce computational time, we downsampled each `fastq` file to 10 millions reads using the sample option in `seqtk` v1.4 (github.com/lh3/seqtk) prior to mapping. We then downsampled read counts to 1,000,000 reads per sample with replacement for 100 replicates and ran `BeXY` with the task `infer` and `SATC`. Because the performance of `SATC` turned out to be rather poor when no depth filter was used, we ran it only on scaffolds that had a normalized depth in the interval (0.3, 2), which corresponds to their default filter Nursyifa et al. (2022). For `BeXY`, we only excluded scaffolds that had zero counts for all samples, as these are not identifiable. We considered all scaffold type classifications by `BeXY` as confident if their state posterior probabilities P(*t*_*c*_|***n***_*r*_) *>* 0.9. `SATC` does not provide an estimate of uncertainty in classifying scaffold types and we assumed all classifications as confident. But as above, we considered all classifications as uncertain if `SATC` threw an error. We counted all scaffolds that were removed by the filter as a uncertain classification.

## 4 Results

### 4.1 Downsampling experiments

#### Sexing for low-depth data

We first investigated the robustness of `BeXY` to low-depth sequencing data of euploid individuals and compared its performance to that of *R*_*x*_ and *R*_*y*_. As shown in Figure 2A, `BeXY` correctly assigned the sex karyotype with high confidence to *>* 97% of all samples even at only 1, 000 reads per sample and to 24% at 100 reads per sample, with a low false discovery rate (FDR) of around 3% at very low depth. The second most powerful methods was *R*_*x*_, which confidently classified a similar fraction of all samples correctly, albeit with a much elevated rate of misclassifications, which results in a FDR of up to 26% at 100 reads per sample. In contrast, *R*_*y*_ confidently classified much fewer samples correctly at an even much higher FDR of up to 55%.

**Figure 2.**
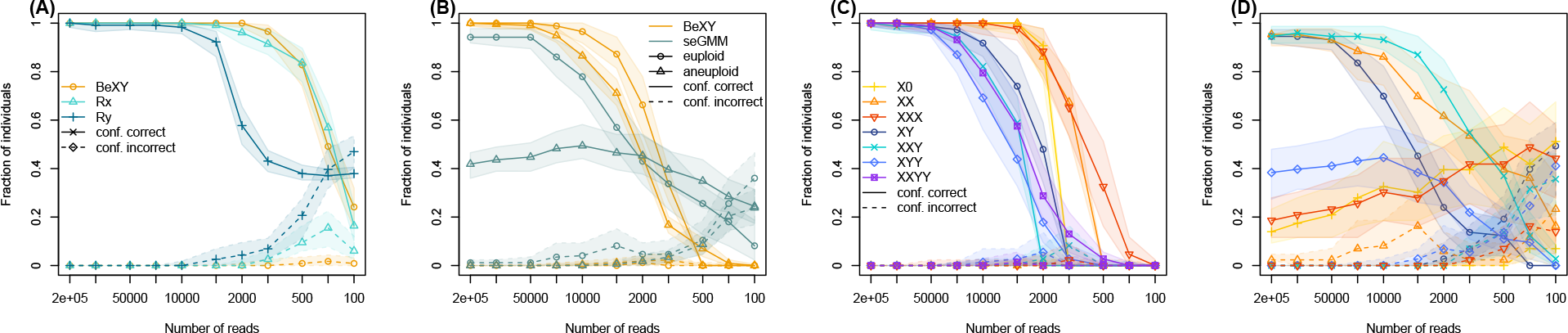
Power to sex individuals. Shown are the median (line with symbols) and 98% confidence interval (shaded area) of the fraction of individuals classified as confidently correct (solid line) and confidently incorrect (dashed line) across 100 downsampling replicates. (A) The performance of `BeXY` compared to *R*_*x*_ and *R*_*y*_ on euploid karyotypes. (B) The power of `BeXY` compared to `seGMM` to classify both euploid and aneuploid karyotypes using a relaxed prior of 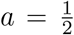. (C) The same results of `BeXY` as in B, but plotted by karyotype. (D) The same results of `seGMM` as in B, but plotted by karyotype.

We then evaluated the power of `BeXY` to detect aneuploid sex karyotypes at low-depth data, and compared it to the competing method `seGMM`. We first evaluated the power of `BeXY` using a relaxed prior on the number of aneuploid individuals of 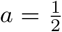. As shown in Figure 2B, `BeXY` correctly assigned both euploid and aneuploid sex karyotypes with high confidence to *>* 90% of all samples at 20,000 reads. We note that the classification accuracy of the euploid karyotypes XY and XX is lower than in Figure 2A, because the prior does not favor euploid karyotypes anymore at noisy counts. `seGMM` correctly assigned *>* 80% of euploid samples and only 47% of aneuploid samples at 20,000 reads. For lower read counts, `seGMM` has a higher power, but also a much elevated FDR of up to 80% for euploid samples and 50% for aneuploid samples.

In Figures 2C and D we show the performance for each karyotype individually. As shown in Figure 2C, the karyotypes that contain a Y chromosome (XY, XXY, XYY and XXYY) require around 20,000 reads to be classified with an accuracy of *>* 80% by `BeXY`, while the karyotypes that contain only X chromosomes (X0, XX and XXX) require only 2,000 reads to obtain the same accuracy. This is explained by the short length of the Y chromosome, which results in small and rather noisy counts. As shown in Figure 2D, classification accuracy of `seGMM` was very variable across karyotypes, with the euploid karyotypes and XXY having a classification accuracy similar to that of `BeXY` and the karyotypes X0, XXX and XYY having low classification accuracy regardless of depth. At low depth, many karyotypes suffered from high misclassification rates, likely because the hard thresholds become unreliable.

When sexing individuals at low depth, the choice of the prior probability on an individual to be aneuploid may have a considerable impact on the performance of `BeXY`. To investigate this we repeated the above inference also with a stringent prior of 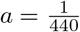, which corresponds to the frequency of aneuploid sex karyotypes in humans and is strongly favoring euploid karyotypes. As a consequence, and in comparison to the case of 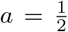, euploid karyotypes required fewer reads (about 1,000) to be classified correctly, while aneuploid karyotypes were wrongly classified as euploids (X0 and XXX as XX and XXY, XYY and XXYY as XY) at low depth (Supplementary Figure 2).

#### Autosomal trisomies for low-depth data

We evaluated the power of `BeXY` to detect autosomal trisomies from low-depth data. As shown in Figure 3, `BeXY` correctly identifies trisomy 21 with high confidence to *>* 96% of all samples at 20, 000 reads per sample and to 47% at 5, 000 reads per sample when using a prior of 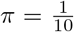. At lower depths, the rate of misclassification increased and samples with trisomy were classified as euploid. `BeXY` correctly assigned autosomal euploidy with a high confidence to 97% of all samples at 10, 000 reads per sample and to 57% at 100 reads per sample. Euploid samples were almost never misclassified as triploid, instead, the uncertainty increased at low depth.

**Figure 3.**
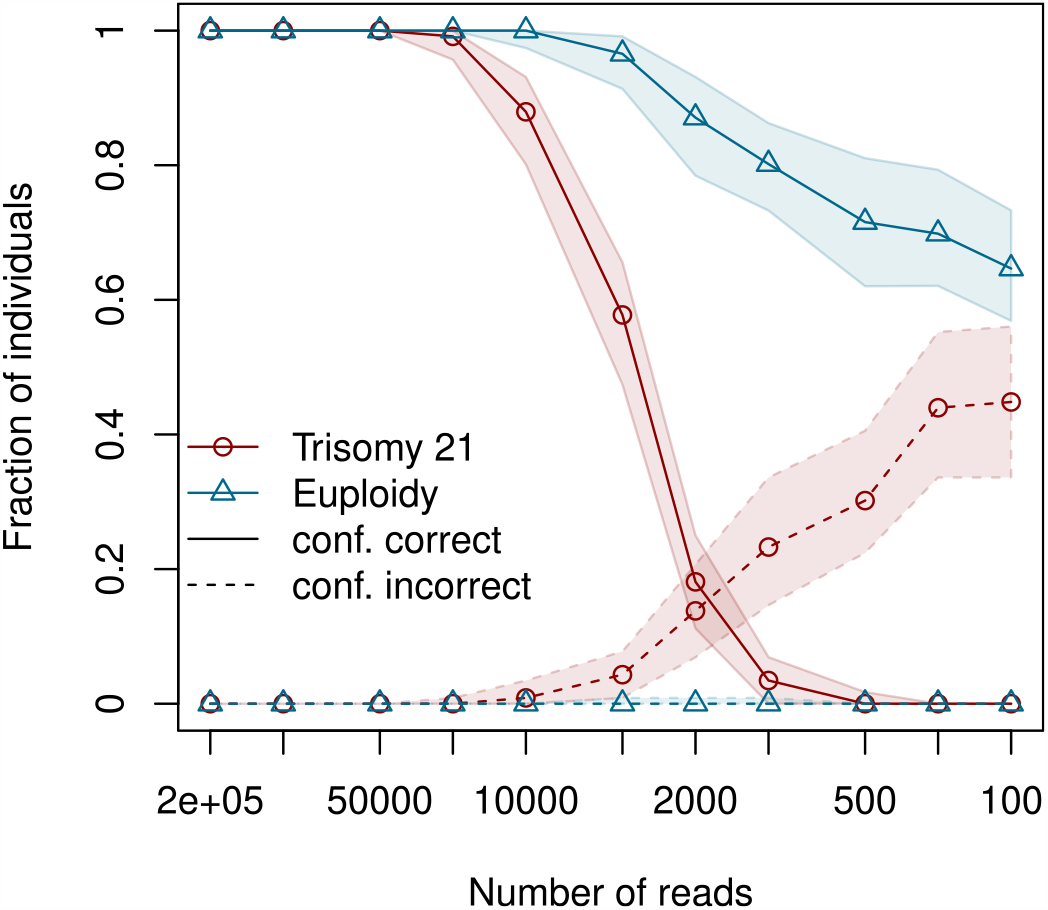
Power to infer autosomal trisomies. The median (line with symbols) and 98% confidence interval (shaded area) of the fraction of individuals classified as confidently correct (solid line) and confidently incorrect (dashed line) as a function of sequencing depth across 100 downsampling replicates when the prior 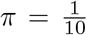. The red line corresponds to the classification of individuals with simulated trisomy 21, while the blue line corresponds to the classification of euploid individuals.

When inferring autosomal trisomies at low depth, the choice of the prior probability on an individual to be triploid may have a considerable impact on the performance of `BeXY`. To investigate this we repeated the above inference also with a more stringent prior of 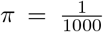, which is strongly favoring euploid autosomes. In this case, euploid samples were almost always correctly classified, even at a depth of only 100 reads per sample, while samples with trisomy 21 required 50, 000 reads to be classified correctly in most cases (98%) and were wrongly classified as euploid at lower depth (Supplementary Figure 3).

#### Joint inference for low-depth data

We next compared the performance of `BeXY` to `SATC` in inferring sex karyotypes jointly with scaffold types. We first investigated the impact of sequencing depth using the full subset of 116 samples. If samples had *>* 1, 000 reads, both methods performed well and correctly assigned the sex karyotype with high confidence to *>* 99% and *>* 95% of all samples respectively (Figure 4A). With lower depth, both methods lose power, but differed in that `BeXY` rarely confidently misclassified individuals while median misclassification rate for `SATC` was up to almost 40%.

**Figure 4.**
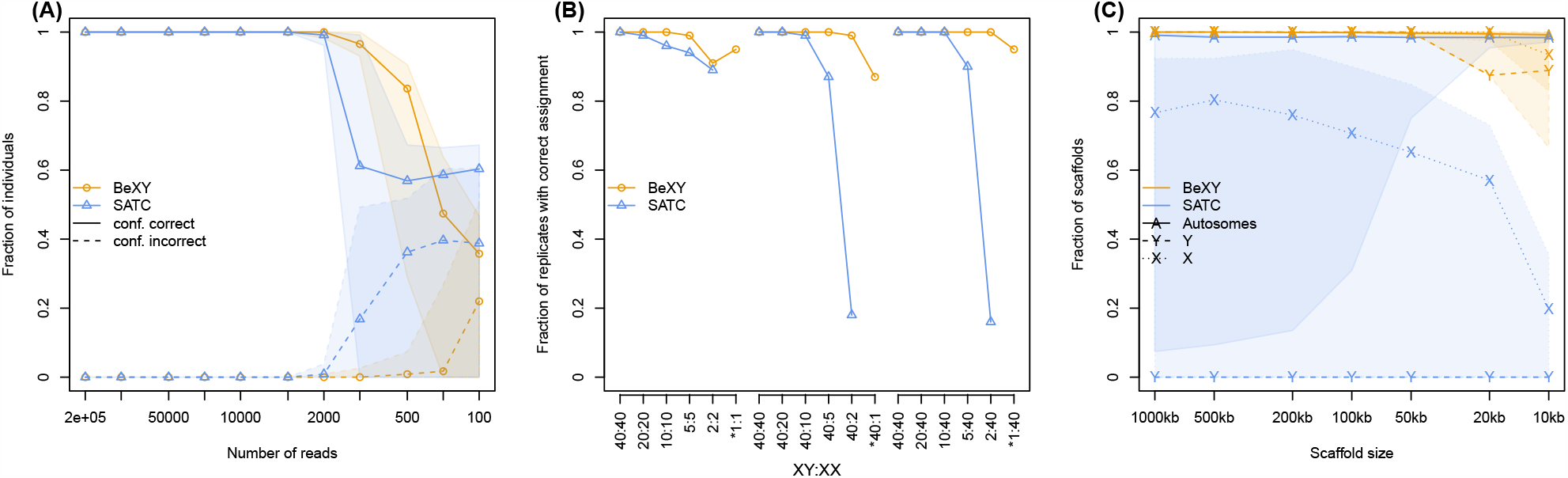
Power to infer sex karyotypes of individuals and scaffold types jointly. (A) The median (line with symbols) and 98% confidence interval (shaded area) of the fraction of individuals classified as confidently correct (solid line) and confidently incorrect (dashed line) as a function of sequencing depth across 100 downsampling replicates for both `BeXY` and `SATC`. (B) The fraction of replicates where the sex karyotype of all individuals was correctly assigned for small (left) and imbalanced (middle and right) data sets. Note that `SATC` requires at least two individuals per karyotype and could not be evaluated for the data sets marked with an asterisk. (C) The power of `BeXY` and `SATC` to classify scaffold types for low-quality reference genomes. Shown are the median (line with symbols) and 98% confidence interval (shaded area) of the fraction of scaffolds classified as confidently correct as a function of scaffold length across 100 replicates for each scaffold type. Note that `SATC` does not classify Y-linked scaffolds.

We next investigated the impact of small sample sizes, imbalanced sex ratios, or both. In 98% and 72% of all simulations conducted, all samples were correctly classified by `BeXY` and `SATC`, respectively. In 27% and 87% of all cases with misclassifications, `BeXY` and `SATC` misclassified all samples to the opposite sex karyotype, suggesting that the challenge at low sample number or with imbalance sex ratios lies in correctly identifying sex-linked scaffolds. As shown in Figure 4B, the fraction of replicates in which all individuals were correctly classified was generally higher for `BeXY` than `SATC`, especially for data set with imbalanced sex ratios. If a sex karyotype was represented by only two individuals, for instance, `SATC` classified all samples correctly in less than 20% of the replicates. In comparison, `BeXY` still classified all samples correctly in 99% of the replicates under these conditions. Notably, `BeXY` also classified all samples correctly in more than 87% of the replicates if a sex karyotype was represented by a single individual - a case in which `SATC` does not attempt any classification.

#### Low-quality reference genome assemblies

We investigated the power of `BeXY` to identify scaffold types in the case of low-quality reference genome assemblies. The simulated reference genome consisted of 10,574 scaffolds ranging from 10^6^ to 10^4^ bases in length. As shown in Figure 4C, the median classification accuracy of `BeXY` was *>* 99% for all larger scaffolds, but dropped slightly to about 93% and 89% for the smallest X-linked and Y-linked scaffolds simulated, respectively. `SATC` had similar power to identify autosomes (median of *>* 98% for all scaffold lengths), although it interestingly misclassified a few long scaffolds as abnormally sex-linked in some replicates. While `SATC` does not classify Y-linked scaffolds, the median classification accuracy for X-linked scaffolds was around 75% for long scaffolds and dropped to 20% for short scaffolds of 10kb, albeit high variability between replicates.

### 4.2 Application to ancient human data set

We inferred sex karyotypes of 954 ancient human samples sequenced with whole-genome shotgun sequencing with the task `infer`. As visualized in Figure 5AC, all individuals were classified with high confidence as XY (*n* = 608) or XX (*n* = 345) except YGS-B2, which was classified as XXY with 100% posterior probability, in line with earlier reports (Ebenesersdóttir et al., 2018). We did not detect any sample with trisomy 21 in this data set.

**Figure 5.**
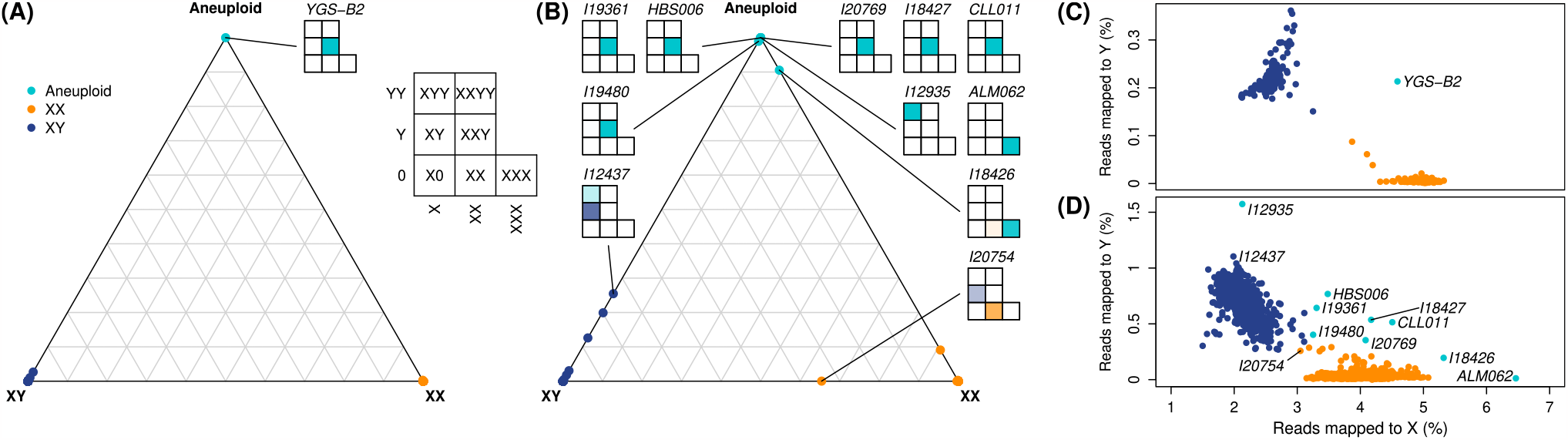
Sex karyotype classification of ancient samples. (A,B) Ternary plot showing the posterior probabilities for each sample to have an XY, XX or aneuploid sex karyotype for the ancient WGS (A) and 1240k target enrichment capture (B) data sets. The color of the points refers to the posterior mode of the sex karyotype. The per karyotype posteriors of all samples identified as aneuploid or with uncertain sex karyotypes are shown using shaded within a 7-cell matrix of the x-axis corresponds to having one, two or three copies of the X-chromosome and the y-axis corresponds to having zero, one or two copies of the Y-chromosome. The samples CLL011.A0101.TF1.1 and ALM062.A0101.TF1.1 were abbreviated to CLL011 and ALM062, respectively. (C,D) Scatter plot for the ancient WGS (C) and 1240k target enrichment capture (D) data sets showing the percentage of reads mapped to Y against the percentage of reads mapped to X, with the same individuals highlighted as in A and B.

We next run the task `infer` on 2,457 ancient human samples sequenced with the 1240k capture target enrichment technique. As visualized in Figure 5BD, we identified 1,141 samples as XX and 1,303 samples as XY with high confidence (posterior probability for the given karyotype larger than 90%). We confirmed the karyotype of all three samples that were previously classified as aneuploid: ALM062.A0101.TF1.1 (XXX, Villalba-Mouco et al., 2021), CLL011.A0101.TF1.1 (XXY, Villalba-Mouco et al., 2021) and HBS006 (XXY, Rivollat et al., 2020) with very high confidence (100% posterior probability for the given karyotype). In addition, we identified six novel aneuploid samples with high confidence: I12935 (XYY, 100%), I18427 (XXY, 100%), I20769 (XXY, 100%), I19361 (XXY, 100%), I19480 (XXY, 99%), and I18426 (XXX, 90.5%). For two samples, the karyotype remains unclear: I12437 had a posterior probability of 25.5% to be XYY and 74.5% to be XY and I20754 had a posterior probability of 34.6% to be XX and 65.4% to be XY. This last sample had the lowest read count of all samples (6,983 reads) and was previously reported as being likely contaminated (Harney et al., 2021), which may explain its uncertain sex karyotype assignment. Finally, we identified a single sample, I19991, of having trisomy 21 with high confidence (99.5%), while all other samples were confidently classified has diploid on chromosome 21 (>99%).

To investigate the effect of the prior choice on aneuploid sex karyotypes, we re-analyzed all individuals with the task `sex` and different values of *a* (Supplementary Figure 4). As expected, the number of individuals classified as aneuploid increases when using less stringent priors. When using the extreme case of 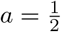, the maximum *a posteriori* estimate was the same sex karyotype for 99.7% and 99.6% of all samples as was obtained with the task `infer` above for the whole-genome and the 1240k capture data set, with an additional three and ten individuals for which an aneuploid sex karyotypes was inferred with high confidence (posterior probability *>* 90%), respectively. However, many of these are likely false positives, suggesting that a more stringent prior should be used. When using 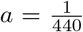 that matches the expected frequency of aneuploid sex karyotypes in humans, the same individuals were classified as aneuploid as when using the task `infer`.

### 4.3 Application to modern human data set

Many factors affect mapping statistics, including how sequencing libraries were generated as well as bioinformatic choices during mapping and filtering of reads. The accuracy of `BeXY` to sex an individual is therefore likely reduced if the parameters ***ρ, γ***_*m*_ and *σ*_*m*_ were inferred on a data set of different characteristics. To explore the sensitivity of `BeXY` to such deviations, we ran `BeXY` on publicly available CRAM-files for 276 modern humans of the Simons Genome Diversity Project (SGDP, Mallick et al., 2016). We first used `BeXY` to sex each sample using the parameters ***ρ, γ***_*m*_ and *σ*_*m*_ inferred on the full set of 954 ancient WGS samples used above. In that case, all but one sample were identified as aneuploidy with high confidence (posterior probability *>* 90%, Supplementary Figure 5AB) with samples reported as females inferred as XXY and samples reported as males as XYY. This seems to be a consequence of the much higher fraction of reads of SGDP samples mapping to the Y-chromosome (around 1.0% for males and 0.15% for females) than of the ancient samples (0.25% for males and almost 0.0% for females, see Figure 5C and Supplementary Figure 5A).

A major difference between the `ATLAS`-pipeline we used to process the ancient samples and the pipeline used by the SGDP is that the former filters out reads with low mapping quality (<30) while the latter does not. Applying the same mapping quality filter on the SGDP CRAM-files leads to a noticeable reduction of reads mapping to the Y chromosome (Supplementary Figure 5C). When sexing individuals with `BeXY` on that data set, only one individual was confidently classified as having an aneuploid sex karyotype (Supplementary Figure 5CD). However, most individuals had very uncertain sex karyotype classifications, suggesting that applying the additional mapping quality filter was not sufficient to correct for the mismatch between the two pipelines.

We next re-estimated all parameters for the SGDP data by running the task `infer` of `BeXY` on the unfiltered CRAM files. That run resulted in most samples being confidently classified to have euploid sex karyotypes (Supplementary Figure 5EF) and only four samples were confidently classified to have aneuploid sex karyotypes: HGDP00725 (X0, 100%), HGDP01036 (XYY, 100%) HGDP01286 (XYY, 100%) and HGDP00546 (XYY, 95.8%). However, SGDP samples were generated with two different sequencing libraries and visual inspection showed that mapping statistics differed noticeably between these groups (Supplementary Figure 5G). Specifically, the PCR-based samples generally had more reads mapping to the Y-chromosome, explaining why `BeXY` classified some of the XY individuals as having an extra Y-chromosome (XYY). We therefore re-ran the task `infer` of `BeXY` while specifying the two library preparation protocols as distinct sequencing types (Supplementary Figure 5GH). In this case, no sample was classified has having trisomy 21 and only a single sample was classified confidently as aneuploid: HGDP00725 (X0, 100%). Although this sample was classified as XX in the original publication (Mallick et al., 2016), our classification as X0 is in line with its heterozygosity on the X chromosome, which was the lowest across all samples (including males). As this sample was sequenced based on a cell line, it is possible that this is an artefact that arose during cell line culturing (e.g. Bergström et al., 2020).

### 4.4 Application to non-model organisms

We ran `BeXY` on the six mammal and bird WGS data sets provided by (Nursyifa et al., 2022) and compared the sex and scaffold-type assignment to that obtained with `SATC`. For all data sets, the two methods did not differ in the sex karyotype classifications. Both methods also mostly agreed in their scaffold-type assignments and classified on average 94.9% of all scaffold identically across data sets. The most common difference in scaffold-type was that `SATC` classified a scaffold as ”abnormally sex-linked“ while `BeXY` classified it as autosomal or X-linked. Notably, and while `SATC` cannot detect Y/W-linked scaffolds, `BeXY` found clear evidence of Y/W-linked scaffolds in muskox, waterbuck and finches, which `SATC` labelled as ”abnormally sex linked“. Specifically, `BeXY` classified the scaffolds 213 and 46825 in muskox, the scaffolds 6207, 8195 11649, 13014 and 16829 in impala, the scaffolds 603, 1604, 2519, 3065, 3205, 3507, 3814, 4043, 4087, 4219, 4352 and 4785 in waterbuck as well as the scaffolds NW_005054686.1, NW_005054701.1, NW_005054702.1, NW_005054723.1, NW_005054728.1, NW_005054736.1, NW_005054744.1, NW_005054750.1, NW_005054752.1, NW_005054753.1, NW_005054754.1 and NW_005054755.1 in finches with a 100% posterior probability as being Y/W-linked.

## 5 Discussion

With the rapid growth of population genomics studies for non-model organisms and ancient samples, there is a need for accurate sex karyotype and scaffold type assignment for low-depth samples. We here present `BeXY`, a Bayesian method that jointly infers sex karyotypes and identifies sex-linked scaffolds from mapping statistics. Our method is flexible with respect to existing knowledge (e.g. scaffold types), retains high accuracy for low-depth samples, is robust to small data sets or those with highly imbalanced sex ratios, is capable of identifying autosomal trisomies, and outperforms all existing methods in the field.

One use case of `BeXY` are non-model organisms, for which both the genetic sex of individuals as well as scaffold types are unknown. `BeXY` only requires mapping statistics, namely the number of reads mapping to each scaffold as well as scaffold lengths, and does not require any prior knowledge on sex karyotypes or sex-linked scaffolds. Importantly, `BeXY` introduces a continuous parametrization of the expected ploidy. This allows for the accommodation of noisy scaffolds showing aberrant ploidy ratios due to mapping issues or mistakes in the assembly, which was previously shown to complicate the identification of sex-linked scaffolds (Nursyifa et al., 2022). Using simulations, we showed that `BeXY` has much higher power to accurately infer scaffold types than the competing methods `SATC` (Nursyifa et al., 2022), particularly for data sets with only few individuals or highly imbalanced sex ratios, and also for low-depth samples (1, 000 reads per sample). In contrast to `SATC`, `BeXY` also accurately identifies Y-linked scaffolds, which `SATC` puts in the same class as those with aberrant ploidy ratios.

The second major use case of `BeXY` is the genetic sexing of (single) individuals, for example for ancient humans. Genetically sexing individuals with `BeXY` has several advantages over existing methods such as *R*_*x*_, *R*_*y*_, `SATC` and `seGMM`. First, `BeXY` is a Bayesian method that calculates the posterior probabilities for each sex karyotype, whereas all existing methods that do sex assignment from read count data are based on hard thresholds. As we showed with simulations, the use of hard thresholds is problematic in the case low-depth samples, which were often un- or misclassified. Second, `BeXY` accurately identifies aneuploid sex karyotypes, which are expected to be found in most large data sets. There exists currently only a single method, `seGMM`, that identifies aneuploid karyotypes from read counts, and as we showed with simulations, `BeXY` has considerably higher power to accurately infer such karyotypes, only requiring about 20,000 reads per individual in the case of ancient human samples.

To sex an individual, `BeXY` only requires mapping statistics of the individual, along with estimates of parameters ***ρ, γ***_*m*_ and *σ*_*m*_ that have been inferred on a larger data set. It is important that these parameters were inferred not only from a data set of the same species, but also from data produced with the same sequencing method, as different methods may differ in their per-scaffold expected read counts. In the case of ancient human samples, for instance, whole-genome shotgun and target enrichment capture data differs considerably in their distribution of reads across chromosomes, as the former is mainly influenced by the length of each chromosome, and the latter mainly by the number of sites targeted per chromosome. However, and as our analyses of modern human samples indicates, more subtle differences in both the preparation of sequencing libraries (e.g. PCR-based vs PCR-free) as well as bioinformatic pipelines (e.g. mapping quality filter) may lead to considerable differences in mapping statistics and hence parameters learned on one data set may not be transferrable to another data set easily and lead to error-prone inference of sex karyotypes. To avoid such issues, we thus recommend to use the task `infer` to learn all relevant parameters if the number of samples permits, i.e. if at least 5-10 samples are at hand. We assume this to be the case for most applications.

Since `BeXY` is a Bayesian method, the choice of the prior may have a big influence if data is limited. That is particularly true for *a*, the expected fraction of individuals with aneuploid sex karyotypes in the sample. Our simulations suggest that if a small value of *a* was used, such as the known frequency of humans with aneuploid sex karyotype 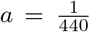, individuals with euploid sex karyotypes should be accurately sexed with as few as 1,000 reads, but individuals with aneuploid sex karyotypes would often be confidently misclassified as euploids as this little data can not overcome the prior. If a larger value of 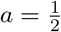 was used, our simulations indicate that both euploids as well as aneuploids individuals should hardly ever be misclassified, but that about 20,000 reads per sample are required for confident classification. The analysis of real data, however, suggested that more reads may be required in practice as we inferred aneuploid sex karyotypes for several ancient human samples with 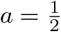 but not with 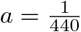. Ultimately, the choice of prior is thus a question of sensitivity versus specificity: If the goal is to ensure that all aneuploid samples have been identified (e.g. to flag them for downstream analysis), a relaxed prior of 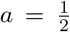 is advised. If, however, the goal is to infer aneuploids with a low false positive rate, a more stringent prior is advised.

## 6 Acknowledgements

This research was funded by a Swiss National Science foundation grant with number 310030_200420 to DW. All computations were done on the IBU cluster at the University of Bern, Switzerland.

## 7 Data Accessibility and Benefit-Sharing

All scripts and data sets (input files) used here are available at https://doi.org/10.5281/zenodo.8344991, where we also provide a complete table of all previously published sequencing data used.

## 8 Author Contributions

The work was conceived and the research designed by MC, CL and DW. The data was analysed by RE and MC. MC and DW wrote the manuscript with input from RE. All authors read and approved the final version of the manuscript.

## References

Allentoft, M. E., Sikora, M., Sjögren, K.-G., Rasmussen, S., Rasmussen, M., Stenderup, J., Damgaard, P. B., Schroeder, H., Ahlström, T., Vinner, L., Malaspinas, A.-S., Margaryan, A., Higham, T., Chivall, D., Lynnerup, N., Harvig, L., Baron, J., Casa, P. D., Dąbrowski, P., Duffy, P. R., Ebel, A. V., Epimakhov, A., Frei, K., Furmanek, M., Gralak, T., Gromov, A., Gronkiewicz, S., Grupe, G., Hajdu, T., Jarysz, R., Khartanovich, V., Khokhlov, A., Kiss, V., Kolář, J., Kriiska, A., Lasak, I., Longhi, C., McGlynn, G., Merkevicius, A., Merkyte, I., Metspalu, M., Mkrtchyan, R., Moiseyev, V., Paja, L., Pálfi, G., Pokutta, D., Pospieszny, Ł., Price, T. D., Saag, L., Sablin, M., Shishlina, N., Smrčka, V., Soenov, V. I., Szeverényi, V., Tóth, G., Trifanova, S. V., Varul, L., Vicze, M., Yepiskoposyan, L., Zhitenev, V., Orlando, L., Sicheritz-Pontén, T., Brunak, S., Nielsen, R., Kristiansen, K., and Willerslev, E. (2015). Population genomics of Bronze Age Eurasia. Nature, 522(7555):167–172.

Amemiya, H. M., Kundaje, A., and Boyle, A. P. (2019). The ENCODE Blacklist: Identification of Problematic Regions of the Genome. Scientific Reports, 9(1):9354.

Antonio, M. L., Gao, Z., Moots, H. M., Lucci, M., Candilio, F., Sawyer, S., Oberreiter, V., Calderon, D., Devitofranceschi, K., Aikens, R. C., Aneli, S., Bartoli, F., Bedini, A., Cheronet, O., Cotter, D. J., Fernandes, D. M., Gasperetti, G., Grifoni, R., Guidi, A., La Pastina, F., Loreti, E., Manacorda, D., Matullo, G., Morretta, S., Nava, A., Fiocchi Nicolai, V., Nomi, F., Pavolini, C., Pentiricci, M., Pergola, P., Piranomonte, M., Schmidt, R., Spinola, G., Sperduti, A., Rubini, M., Bondioli, L., Coppa, A., Pinhasi, R., and Pritchard, J. K. (2019). Ancient Rome: A genetic crossroads of Europe and the Mediterranean. Science, 366(6466):708–714.

Bachtrog, D., Mank, J. E., Peichel, C. L., Kirkpatrick, M., Otto, S. P., Ashman, T.-L., Hahn, M. W., Kitano, J., Mayrose, I., Ming, R., Perrin, N., Ross, L., Valenzuela, N., Vamosi, J. C., and The Tree of Sex Consortium (2014). Sex Determination: Why So Many Ways of Doing It? PLoS Biology, 12(7):e1001899.

Benestan, L., Moore, J.-S., Sutherland, B. J. G., Le Luyer, J., Maaroufi, H., Rougeux, C., Normandeau, E., Rycroft, N., Atema, J., Harris, L. N., Tallman, R. F., Greenwood, S. J., Clark, F. K., and Bernatchez, L. (2017). Sex matters in massive parallel sequencing: Evidence for biases in genetic parameter estimation and investigation of sex determination systems. Molecular Ecology, 26(24):6767–6783.

Bergström, A., McCarthy, S. A., Hui, R., Almarri, M. A., Ayub, Q., Danecek, P., Chen, Y., Felkel, S., Hallast, P., Kamm, J., Blanché, H., Deleuze, J.-F., Cann, H., Mallick, S., Reich, D., Sandhu, M. S., Skoglund, P., Scally, A., Xue, Y., Durbin, R., and Tyler-Smith, C. (2020). Insights into human genetic variation and population history from 929 diverse genomes. Science, 367(6484):eaay5012.

Bidon, T., Janke, A., Fain, S. R., Eiken, H. G., Hagen, S. B., Saarma, U., Hallström, B. M., Lecomte, N., and Hailer, F. (2014). Brown and Polar Bear Y Chromosomes Reveal Extensive Male-Biased Gene Flow within Brother Lineages. Molecular Biology and Evolution, 31(6):1353–1363.

Breman, A. and Stankiewicz, P. (2021). Karyotyping as the first genomic approach. In Genomics of Rare Diseases: Understanding Disease Genetics Using Genomic Approaches, number 17-31. Academic Press, an imprint of Elsevier, London, United Kingdom ; San Diego, CA.

Broushaki, F., Thomas, M. G., Link, V., López, S., Van Dorp, L., Kirsanow, K., Hofmanová, Z., Diekmann, Y., Cassidy, L. M., Díez-del-Molino, D., Kousathanas, A., Sell, C., Robson, H. K., Martiniano, R., Blöcher, J., Scheu, A., Kreutzer, S., Bollongino, R., Bobo, D., Davoudi, H., Munoz, O., Currat, M., Abdi, K., Biglari, F., Craig, O. E., Bradley, D. G., Shennan, S., Veeramah, K. R., Mashkour, M., Wegmann, D., Hellenthal, G., and Burger, J. (2016). Early Neolithic genomes from the eastern Fertile Crescent. Science, 353(6298):499–503.

Buonasera, T., Eerkens, J., De Flamingh, A., Engbring, L., Yip, J., Li, H., Haas, R., DiGiuseppe, D., Grant, D., Salemi, M., Nijmeh, C., Arellano, M., Leventhal, A., Phinney, B., Byrd, B. F., Malhi, R. S., and Parker, G. (2020). A comparison of proteomic, genomic, and osteological methods of archaeological sex estimation. Scientific Reports, 10(1):11897.

Burger, J., Link, V., Blöcher, J., Schulz, A., Sell, C., Pochon, Z., Diekmann, Y., Žegarac, A., Hofmanová, Z., Winkelbach, L., Reyna-Blanco, C. S., Bieker, V., Orschiedt, J., Brinker, U., Scheu, A., Leuenberger, C., Bertino, T. S., Bollongino, R., Lidke, G., Stefanović, S., Jantzen, D., Kaiser, E., Terberger, T., Thomas, M. G., Veeramah, K. R., and Wegmann, D. (2020). Low Prevalence of Lactase Persistence in Bronze Age Europe Indicates Ongoing Strong Selection over the Last 3,000 Years. Current Biology, 30(21):4307–4315.e13.

Cabrera, A. A., Rey-Iglesia, A., Louis, M., Skovrind, M., Westbury, M. V., and Lorenzen, E. D. (2022). How low can you go? Introducing SeXY: Sex identification from low-quantity sequencing data despite lacking assembled sex chromosomes. Ecology and Evolution, 12(8).

Damgaard, P. D. B., Marchi, N., Rasmussen, S., Peyrot, M., Renaud, G., Korneliussen, T., Moreno-Mayar, J. V., Pedersen, M. W., Goldberg, A., Usmanova, E., Baimukhanov, N., Loman, V., Hedeager, L., Pedersen, A. G., Nielsen, K., Afanasiev, G., Akmatov, K., Aldashev, A., Alpaslan, A., Baimbetov, G., Bazaliiskii, V. I., Beisenov, A., Boldbaatar, B., Boldgiv, B., Dorzhu, C., Ellingvag, S., Erdenebaatar, D., Dajani, R., Dmitriev, E., Evdokimov, V., Frei, K. M., Gromov, A., Goryachev, A., Hakonarson, H., Hegay, T., Khachatryan, Z., Khaskhanov, R., Kitov, E., Kolbina, A., Kubatbek, T., Kukushkin, A., Kukushkin, I., Lau, N., Margaryan, A., Merkyte, I., Mertz, I. V., Mertz, V. K., Mijiddorj, E., Moiyesev, V., Mukhtarova, G., Nurmukhanbetov, B., Orozbekova, Z., Panyushkina, I., Pieta, K., Smrčka, V., Shevnina, I., Logvin, A., Sjögren, K.-G., Štolcová, T., Taravella, A. M., Tashbaeva, K., Tkachev, A., Tulegenov, T., Voyakin, D., Yepiskoposyan, L., Undrakhbold, S., Varfolomeev, V., Weber, A., Wilson Sayres, M. A., Kradin, N., Allentoft, M. E., Orlando, L., Nielsen, R., Sikora, M., Heyer, E., Kristiansen, K., and Willerslev, E. (2018). 137 ancient human genomes from across the Eurasian steppes. Nature, 557(7705):369–374.

De Barros Damgaard, P., Martiniano, R., Kamm, J., Moreno-Mayar, J. V., Kroonen, G., Peyrot, M., Barjamovic, G., Rasmussen, S., Zacho, C., Baimukhanov, N., Zaibert, V., Merz, V., Biddanda, A., Merz, I., Loman, V., Evdokimov, V., Usmanova, E., Hemphill, B., Seguin-Orlando, A., Yediay, F. E., Ullah, I., Sjögren, K.-G., Iversen, K. H., Choin, J., De La Fuente, C., Ilardo, M., Schroeder, H., Moiseyev, V., Gromov, A., Polyakov, A., Omura, S., Senyurt, S. Y., Ahmad, H., McKenzie, C., Margaryan, A., Hameed, A., Samad, A., Gul, N., Khokhar, M. H., Goriunova, O. I., Bazaliiskii, V. I., Novembre, J., Weber, A. W., Orlando, L., Allentoft, M. E., Nielsen, R., Kristiansen, K., Sikora, M., Outram, A. K., Durbin, R., and Willerslev, E. (2018). The first horse herders and the impact of early Bronze Age steppe expansions into Asia. Science, 360(6396):eaar7711.

De La Cruz, I., González-Oliver, A., Kemp, B. M., Román, J. A., Smith, D. G., and Torre-Blanco, A. (2008). Sex Identification of Children Sacrificed to the Ancient Aztec Rain Gods in Tlatelolco. Current Anthropology, 49(3):519–526.

Ebenesersdóttir, S. S., Sandoval-Velasco, M., Gunnarsdóttir, E. D., Jagadeesan, A., Guðmundsdóttir, V. B., Thordardóttir, E. L., Einarsdóttir, M. S., Moore, K. H. S., Sigurðsson, Á., Magnúsdóttir, D. N., Jónsson, H., Snorradóttir, S., Hovig, E., Møller, P., Kockum, I., Olsson, T., Alfredsson, L., Hansen, T. F., Werge, T., Cavalleri, G. L., Gilbert, E., Lalueza-Fox, C., Walser, J. W., Kristjánsdóttir, S., Gopalakrishnan, S., Árnadóttir, L., Magnússon, Ó. Þ., Gilbert, M. T. P., Stefánsson, K., and Helgason, A. (2018). Ancient genomes from Iceland reveal the making of a human population. Science, 360(6392):1028–1032.

Ellegren, H. (2007). Characteristics, causes and evolutionary consequences of male-biased mutation. Proceedings of the Royal Society B: Biological Sciences, 274(1606):1–10.

Ellegren, H. (2009). The different levels of genetic diversity in sex chromosomes and autosomes. Trends in Genetics, 25(6):278–284.

Ellegren, H. (2014). Genome sequencing and population genomics in non-model organisms. Trends in Ecology & Evolution, 29(1):51–63.

Fairbairn, D. J., Blanckenhorn, Wolf U., and Tamás, S. (2007). Sex, Size and Gender Roles: Evolutionary Studies of Sexual Size Dimorphism. Oxford University Press (OUP).

Fernandes, D. M., Mittnik, A., Olalde, I., Lazaridis, I., Cheronet, O., Rohland, N., Mallick, S., Bernardos, R., Broomandkhoshbacht, N., Carlsson, J., Culleton, B. J., Ferry, M., Gamarra, B., Lari, M., Mah, M., Michel, M., Modi, A., Novak, M., Oppenheimer, J., Sirak, K. A., Stewardson, K., Mandl, K., Schattke, C., Özdoğan, K. T., Lucci, M., Gasperetti, G., Candilio, F., Salis, G., Vai, S., Camarós, E., Calò, C., Catalano, G., Cueto, M., Forgia, V., Lozano, M., Marini, E., Micheletti, M., Miccichè, R. M., Palombo, M. R., Ramis, D., Schimmenti, V., Sureda, P., Teira, L., Teschler-Nicola, M., Kennett, D. J., Lalueza-Fox, C., Patterson, N., Sineo, L., Coppa, A., Caramelli, D., Pinhasi, R., and Reich, D. (2020). The spread of steppe and Iranian-related ancestry in the islands of the western Mediterranean. Nature Ecology & Evolution, 4(3):334–345.

Fowler, C., Olalde, I., Cummings, V., Armit, I., Büster, L., Cuthbert, S., Rohland, N., Cheronet, O., Pinhasi, R., and Reich, D. (2022). A high-resolution picture of kinship practices in an Early Neolithic tomb. Nature, 601(7894):584–587.

Fu, Q., Posth, C., Hajdinjak, M., Petr, M., Mallick, S., Fernandes, D., Furtwängler, A., Haak, W., Meyer, M., Mittnik, A., Nickel, B., Peltzer, A., Rohland, N., Slon, V., Talamo, S., Lazaridis, I., Lipson, M., Mathieson, I., Schiffels, S., Skoglund, P., Derevianko, A. P., Drozdov, N., Slavinsky, V., Tsybankov, A., Cremonesi, R. G., Mallegni, F., Gély, B., Vacca, E., Morales, M. R. G., Straus, L. G., Neugebauer-Maresch, C., Teschler-Nicola, M., Constantin, S., Moldovan, O. T., Benazzi, S., Peresani, M., Coppola, D., Lari, M., Ricci, S., Ronchitelli, A., Valentin, F., Thevenet, C., Wehrberger, K., Grigorescu, D., Rougier, H., Crevecoeur, I., Flas, D., Semal, P., Mannino, M. A., Cupillard, C., Bocherens, H., Conard, N. J., Harvati, K., Moiseyev, V., Drucker, D. G., Svoboda, J., Richards, M. P., Caramelli, D., Pinhasi, R., Kelso, J., Patterson, N., Krause, J., Pääbo, S., and Reich, D. (2016). The genetic history of Ice Age Europe. Nature, 534(7606):200–205.

Furtwängler, A., Rohrlach, A. B., Lamnidis, T. C., Papac, L., Neumann, G. U., Siebke, I., Reiter, E., Steuri, N., Hald, J., Denaire, A., Schnitzler, B., Wahl, J., Ramstein, M., Schuenemann, V. J., Stockhammer, P. W., Hafner, A., Lösch, S., Haak, W., Schiffels, S., and Krause, J. (2020). Ancient genomes reveal social and genetic structure of Late Neolithic Switzerland. Nature Communications, 11(1):1915.

Gamba, C., Jones, E. R., Teasdale, M. D., McLaughlin, R. L., Gonzalez-Fortes, G., Mattiangeli, V., Domboróczki, L., Kővári, I., Pap, I., Anders, A., Whittle, A., Dani, J., Raczky, P., Higham, T. F. G., Hofreiter, M., Bradley, D. G., and Pinhasi, R. (2014). Genome flux and stasis in a five millennium transect of European prehistory. Nature Communications, 5(1):5257.

Gao, F., Chang, D., Biddanda, A., Ma, L., Guo, Y., Zhou, Z., and Keinan, A. (2015). XWAS: A Software Toolset for Genetic Data Analysis and Association Studies of the X Chromosome. Journal of Heredity, 106(5):666–671.

Gautier, M. (2014). Using genotyping data to assign markers to their chromosome type and to infer the sex of individuals: A Bayesian model-based classifier. Molecular Ecology Resources, 14(6):1141–1159.

Gower, G., Fenderson, L. E., Salis, A. T., Helgen, K. M., Van Loenen, A. L., Heiniger, H., Hofman-Kamińska, E., Kowalczyk, R., Mitchell, K. J., Llamas, B., and Cooper, A. (2019). Widespread male sex bias in mammal fossil and museum collections. Proceedings of the National Academy of Sciences, 116(38):19019–19024.

Grabherr, M. G., Russell, P., Meyer, M., Mauceli, E., Alföldi, J., Di Palma, F., and Lindblad-Toh, K. (2010). Genome-wide synteny through highly sensitive sequence alignment: Satsuma. Bioinformatics, 26(9):1145–1151.

Grath, S. and Parsch, J. (2016). Sex-Biased Gene Expression. Annual Review of Genetics, 50(1):29–44.

Graves, J. A. M. (2008). Weird Animal Genomes and the Evolution of Vertebrate Sex and Sex Chromosomes. Annual Review of Genetics, 42(1):565–586.

Günther, T., Malmström, H., Svensson, E. M., Omrak, A., Sánchez-Quinto, F., Kılınç, G. M., Krzewińska, M., Eriksson, G., Fraser, M., Edlund, H., Munters, A. R., Coutinho, A., Simões, L. G., Vicente, M., Sjölander, A., Jansen Sellevold, B., Jørgensen, R., Claes, P., Shriver, M. D., Valdiosera, C., Netea, M. G., Apel, J., Lidén, K., Skar, B., Storå, J., Götherström, A., and Jakobsson, M. (2018). Population genomics of Mesolithic Scandinavia: Investigating early postglacial migration routes and high-latitude adaptation. PLOS Biology, 16(1):e2003703.

Harney, É., Cheronet, O., Fernandes, D. M., Sirak, K., Mah, M., Bernardos, R., Adamski, N., Broomandkhoshbacht, N., Callan, K., Lawson, A. M., Oppenheimer, J., Stewardson, K., Zalzala, F., Anders, A., Candilio, F., Constantinescu, M., Coppa, A., Ciobanu, I., Dani, J., Gallina, Z., Genchi, F., Nagy, E. G., Hajdu, T., Hellebrandt, M., Horváth, A., Király, Á., Kiss, K., Kolozsi, B., Kovács, P., Köhler, K., Lucci, M., Pap, I., Popovici, S., Raczky, P., Simalcsik, A., Szeniczey, T., Vasilyev, S., Virag, C., Rohland, N., Reich, D., and Pinhasi, R. (2021). A minimally destructive protocol for DNA extraction from ancient teeth. Genome Research, 31(3):472–483.

Harney, É., Olalde, I., Bruwelheide, K., Barca, K. G., Curry, R., Comer, E., Rohland, N., Owsley, D., and Reich, D. (2022). Technical Report on Ancient DNA analysis of 27 African Americans from Catoctin Furnace, Maryland. Preprint, Genomics.

Hedenstierna-Jonson, C., Kjellström, A., Zachrisson, T., Krzewińska, M., Sobrado, V., Price, N., Günther, T., Jakobsson, M., Götherström, A., and Storå, J. (2017). A female Viking warrior confirmed by genomics. American Journal of Physical Anthropology, 164(4):853–860.

Herrero, A., Klütsch, C. F. C., Holmala, K., Maduna, S. N., Kopatz, A., Eiken, H. G., and Hagen, S. B. (2021). Genetic analysis indicates spatial-dependent patterns of sex-biased dispersal in Eurasian lynx in Finland. PLOS ONE, 16(2):e0246833.

Hofmanová, Z., Kreutzer, S., Hellenthal, G., Sell, C., Diekmann, Y., Díez-del-Molino, D., Van Dorp, L., López, S., Kousathanas, A., Link, V., Kirsanow, K., Cassidy, L. M., Martiniano, R., Strobel, M., Scheu, A., Kotsakis, K., Halstead, P., Triantaphyllou, S., Kyparissi-Apostolika, N., Urem-Kotsou, D., Ziota, C., Adaktylou, F., Gopalan, S., Bobo, D. M., Winkelbach, L., Blöcher, J., Unterländer, M., Leuenberger, C., Çilingiroğlu, Ç., Horejs, B., Gerritsen, F., Shennan, S. J., Bradley, D. G., Currat, M., Veeramah, K. R., Wegmann, D., Thomas, M. G., Papageorgopoulou, C., and Burger, J. (2016). Early farmers from across Europe directly descended from Neolithic Aegeans. Proceedings of the National Academy of Sciences, 113(25):6886–6891.

Johnson, N. L., Kotz, S., and Balakrishnan, N. (1997). Discrete Multivariate Distributions. Wiley Series in Probability and Statistics. Wiley.

Jones, E. R., Gonzalez-Fortes, G., Connell, S., Siska, V., Eriksson, A., Martiniano, R., McLaughlin, R. L., Gallego Llorente, M., Cassidy, L. M., Gamba, C., Meshveliani, T., Bar-Yosef, O., Müller, W., Belfer-Cohen, A., Matskevich, Z., Jakeli, N., Higham, T. F. G., Currat, M., Lordkipanidze, D., Hofreiter, M., Manica, A., Pinhasi, R., and Bradley, D. G. (2015). Upper Palaeolithic genomes reveal deep roots of modern Eurasians. Nature Communications, 6(1):8912.

Jones, E. R., Zarina, G., Moiseyev, V., Lightfoot, E., Nigst, P. R., Manica, A., Pinhasi, R., and Bradley, D. G. (2017). The Neolithic Transition in the Baltic Was Not Driven by Admixture with Early European Farmers. Current Biology, 27(4):576–582.

Kennett, D. J., Lipson, M., Prufer, K. M., Mora-Marín, D., George, R. J., Rohland, N., Robinson, M., Trask, W. R., Edgar, H. H. J., Hill, E. C., Ray, E. E., Lynch, P., Moes, E., O’Donnell, L., Harper, T. K., Kate, E. J., Ramos, J., Morris, J., Gutierrez, S. M., Ryan, T. M., Culleton, B. J., Awe, J. J., and Reich, D. (2022). South-to-north migration preceded the advent of intensive farming in the Maya region. Nature Communications, 13(1):1530.

Kocijan, I., Dolenec, P., Pavokovic, G., and Dolenec, Z. (2011). Sex-typing bird species with little or no sexual dimorphism: An evaluation of molecular and morphological sexing. Journal of Biological Research, 15:145.

Lazaridis, I., Alpaslan-Roodenberg, S., Andreeva, D., Andrija, G., Badalyan, R., Bakardzhiev, S., Balen, J., Bejko, L., Bernardos, R., Bertsatos, A., Biber, H., Bilir, A., Bodru, M., Callan, K., Candilio, F., Cari, M., Ciobanu, I., Demcenco, T. I., Dergachev, V., Derin, Z., Deskaj, S., Devejyan, S., Djordjevi, V., Eccles, L. R., Elenski, N., Engin, A., Erdo, N., Frînculeasa, A., Galaty, M. L., Gamarra, B., Gasparyan, B., Gaydarska, B., Genç, E., Gültekin, T., Gündüz, S., Hajdu, T., Heyd, V., Hobosyan, S., Hovhannisyan, N., Iliev, I., Iliev, L., Iliev, S., Jovanova, L., Karkanas, P., Kavaz-K, B., Khudaverdyan, A., Kiss, K., Krenz-Niedba, M., Levy, T. E., Liritzis, I., Lorentz, K. O., Martirosyan-Olshansky, K., Matthews, R., Matthews, W., McSweeney, K., Melikyan, V., Micco, A., Michel, M., Mila, L., Olalde, I., Oppenheimer, J., Osterholtz, A., Özdemir, C., Özdo, K. T., Papakonstantinou, N., Papathanasiou, A., Paraman, L., Paskary, E. G., Patterson, N., Petrakiev, I., Petrosyan, L., Petrova, V., Philippa-Touchais, A., Piliposyan, A., Kuzman, N. P., Potrebica, H., Preda-B, B., Price, T. D., Qiu, L., Richardson, A., Roodenberg, J., Ruka, R., Russeva, V., Schepartz, L., Selçuk, T., Sevim-Erol, A., Shamoon-Pour, M., Shephard, H. M., Sideris, A., Simalcsik, A., Simonyan, H., Sinika, V., Sirak, K., Sirbu, G., Sönmez-Sözer, Ç., Stathi, M., Steskal, M., Stewardson, K., Stocker, S., Suata-Alpaslan, F., Suvorov, A., Szécsényi-Nagy, A., Szeniczey, T., Telnov, N., Temov, S., Todorova, N., Tota, U., Touchais, G., Triantaphyllou, S., Türker, A., Ugarkovi, M., Walsh, S., Zalzala, F., Zettl, A., Zhang, Z., and Çavu, R. (2022). Ancient DNA from Mesopotamia suggests distinct Pre-Pottery and Pottery Neolithic migrations into Anatolia. Science, 377:982–987.

Lazaridis, I., Mittnik, A., Patterson, N., Mallick, S., Rohland, N., Pfrengle, S., Furtwängler, A., Peltzer, A., Posth, C., Vasilakis, A., McGeorge, P. J. P., Konsolaki-Yannopoulou, E., Korres, G., Martlew, H., Michalodimitrakis, M., Özsait, M., Özsait, N., Papathanasiou, A., Richards, M., Roodenberg, S. A., Tzedakis, Y., Arnott, R., Fernandes, D. M., Hughey, J. R., Lotakis, D. M., Navas, P. A., Maniatis, Y., Stamatoyannopoulos, J. A., Stewardson, K., Stockhammer, P., Pinhasi, R., Reich, D., Krause, J., and Stamatoyannopoulos, G. (2017). Genetic origins of the Minoans and Mycenaeans. Nature, 548(7666):214–218.

Lazaridis, I., Nadel, D., Rollefson, G., Merrett, D. C., Rohland, N., Mallick, S., Fernandes, D., Novak, M., Gamarra, B., Sirak, K., Connell, S., Stewardson, K., Harney, E., Fu, Q., Gonzalez-Fortes, G., Jones, E. R., Roodenberg, S. A., Lengyel, G., Bocquentin, F., Gasparian, B., Monge, J. M., Gregg, M., Eshed, V., Mizrahi, A.-S., Meiklejohn, C., Gerritsen, F., Bejenaru, L., Blüher, M., Campbell, A., Cavalleri, G., Comas, D., Froguel, P., Gilbert, E., Kerr, S. M., Kovacs, P., Krause, J., McGettigan, D., Merrigan, M., Merriwether, D. A., O’Reilly, S., Richards, M. B., Semino, O., Shamoon-Pour, M., Stefanescu, G., Stumvoll, M., Tönjes, A., Torroni, A., Wilson, J. F., Yengo, L., Hovhannisyan, N. A., Patterson, N., Pinhasi, R., and Reich, D. (2016). Genomic insights into the origin of farming in the ancient Near East. Nature, 536(7617):419–424.

Lazaridis, I., Patterson, N., Mittnik, A., Renaud, G., Mallick, S., Kirsanow, K., Sudmant, P. H., Schraiber, J. G., Castellano, S., Lipson, M., Berger, B., Economou, C., Bollongino, R., Fu, Q., Bos, K. I., Nordenfelt, S., Li, H., De Filippo, C., Prüfer, K., Sawyer, S., Posth, C., Haak, W., Hallgren, F., Fornander, E., Rohland, N., Delsate, D., Francken, M., Guinet, J.-M., Wahl, J., Ayodo, G., Babiker, H. A., Bailliet, G., Balanovska, E., Balanovsky, O., Barrantes, R., Bedoya, G., Ben-Ami, H., Bene, J., Berrada, F., Bravi, C. M., Brisighelli, F., Busby, G. B. J., Cali, F., Churnosov, M., Cole, D. E. C., Corach, D., Damba, L., Van Driem, G., Dryomov, S., Dugoujon, J.-M., Fedorova, S. A., Gallego Romero, I., Gubina, M., Hammer, M., Henn, B. M., Hervig, T., Hodoglugil, U., Jha, A. R., Karachanak-Yankova, S., Khusainova, R., Khusnutdinova, E., Kittles, R., Kivisild, T., Klitz, W., Kučinskas, V., Kushniarevich, A., Laredj, L., Litvinov, S., Loukidis, T., Mahley, R. W., Melegh, B., Metspalu, E., Molina, J., Mountain, J., Näkkäläjärvi, K., Nesheva, D., Nyambo, T., Osipova, L., Parik, J., Platonov, F., Posukh, O., Romano, V., Rothhammer, F., Rudan, I., Ruizbakiev, R., Sahakyan, H., Sajantila, A., Salas, A., Starikovskaya, E. B., Tarekegn, A., Toncheva, D., Turdikulova, S., Uktveryte, I., Utevska, O., Vasquez, R., Villena, M., Voevoda, M., Winkler, C. A., Yepiskoposyan, L., Zalloua, P., Zemunik, T., Cooper, A., Capelli, C., Thomas, M. G., Ruiz-Linares, A., Tishkoff, S. A., Singh, L., Thangaraj, K., Villems, R., Comas, D., Sukernik, R., Metspalu, M., Meyer, M., Eichler, E. E., Burger, J., Slatkin, M., Pääbo, S., Kelso, J., Reich, D., and Krause, J. (2014). Ancient human genomes suggest three ancestral populations for present-day Europeans. Nature, 513(7518):409–413.

Li, H. (2013). Aligning sequence reads, clone sequences and assembly contigs with BWA-MEM. arXiv preprint.

Li, H., Handsaker, B., Wysoker, A., Fennell, T., Ruan, J., Homer, N., Marth, G., Abecasis, G., Durbin, R., and 1000 Genome Project Data Processing Subgroup (2009). The Sequence Alignment/Map format and SAMtools. Bioinformatics, 25(16):2078–2079.

Link, V., Kousathanas, A., Veeramah, K., Sell, C., Scheu, A., and Wegmann, D. (2017). ATLAS: Analysis Tools for Low-depth and Ancient Samples. Preprint, Bioinformatics.

Lipson, M., Sawchuk, E. A., Thompson, J. C., Oppenheimer, J., Tryon, C. A., Ranhorn, K. L., De Luna, K. M., Sirak, K. A., Olalde, I., Ambrose, S. H., Arthur, J. W., Arthur, K. J. W., Ayodo, G., Bertacchi, A., Cerezo-Román, J. I., Culleton, B. J., Curtis, M. C., Davis, J., Gidna, A. O., Hanson, A., Kaliba, P., Katongo, M., Kwekason, A., Laird, M. F., Lewis, J., Mabulla, A. Z. P., Mapemba, F., Morris, A., Mudenda, G., Mwafulirwa, R., Mwangomba, D., Ndiema, E., Ogola, C., Schilt, F., Willoughby, P. R., Wright, D. K., Zipkin, A., Pinhasi, R., Kennett, D. J., Manthi, F. K., Rohland, N., Patterson, N., Reich, D., and Prendergast, M. E. (2022). Ancient DNA and deep population structure in sub-Saharan African foragers. Nature, 603(7900):290–296.

Lipson, M., Szécsényi-Nagy, A., Mallick, S., Pósa, A., Stégmár, B., Keerl, V., Rohland, N., Stewardson, K., Ferry, M., Michel, M., Oppenheimer, J., Broomandkhoshbacht, N., Harney, E., Nordenfelt, S., Llamas, B., Gusztáv Mende, B., Köhler, K., Oross, K., Bondár, M., Marton, T., Osztás, A., Jakucs, J., Paluch, T., Horváth, F., Csengeri, P., Koós, J., Sebők, K., Anders, A., Raczky, P., Regenye, J., Barna, J. P., Fábián, S., Serlegi, G., Toldi, Z., Gyöngyvér Nagy, E., Dani, J., Molnár, E., Pálfi, G., Márk, L., Melegh, B., Bánfai, Z., Domboróczki, L., Fernández-Eraso, J., Antonio Mujika-Alustiza, J., Alonso Fernández, C., Jiménez Echevarría, J., Bollongino, R., Orschiedt, J., Schierhold, K., Meller, H., Cooper, A., Burger, J., Bánffy, E., Alt, K. W., Lalueza-Fox, C., Haak, W., and Reich, D. (2017). Parallel palaeogenomic transects reveal complex genetic history of early European farmers. Nature, 551(7680):368–372.

Liu, S., Zeng, Y., Wang, C., Zhang, Q., Chen, M., Wang, X., Wang, L., Lu, Y., Guo, H., and Bu, F. (2022a). seGMM: A New Tool for Gender Determination From Massively Parallel Sequencing Data. Frontiers in Genetics, 13:850804.

Liu, Y.-C., Hunter-Anderson, R., Cheronet, O., Eakin, J., Camacho, F., Pietrusewsky, M., Rohland, N., Ioannidis, A., Athens, J. S., Douglas, M. T., Ikehara-Quebral, R. M., Bernardos, R., Culleton, B. J., Mah, M., Adamski, N., Broomandkhoshbacht, N., Callan, K., Lawson, A. M., Mandl, K., Michel, M., Oppenheimer, J., Stewardson, K., Zalzala, F., Kidd, K., Kidd, J., Schurr, T. G., Auckland, K., Hill, A. V. S., Mentzer, A. J., Quinto-Cortés, C. D., Robson, K., Kennett, D. J., Patterson, N., Bustamante, C. D., Moreno-Estrada, A., Spriggs, M., Vilar, M., Lipson, M., Pinhasi, R., and Reich, D. (2022b). Ancient DNA reveals five streams of migration into Micronesia and matrilocality in early Pacific seafarers. Science, 377(6601):72–79.

Louis, M., Skovrind, M., Garde, E., Heide-Jørgensen, M. P., Szpak, P., and Lorenzen, E. D. (2021). Population-specific sex and size variation in long-term foraging ecology of belugas and narwhals. Royal Society Open Science, 8(2):rsos.202226, 202226.

Madel, M.-B., Niederstätter, H., and Parson, W. (2016). TriXY—Homogeneous genetic sexing of highly degraded forensic samples including hair shafts. Forensic Science International: Genetics, 25:166–174.

Mallick, S., Li, H., Lipson, M., Mathieson, I., Gymrek, M., Racimo, F., Zhao, M., Chennagiri, N., Nordenfelt, S., Tandon, A., Skoglund, P., Lazaridis, I., Sankararaman, S., Fu, Q., Rohland, N., Renaud, G., Erlich, Y., Willems, T., Gallo, C., Spence, J. P., Song, Y. S., Poletti, G., Balloux, F., Van Driem, G., De Knijff, P., Romero, I. G., Jha, A. R., Behar, D. M., Bravi, C. M., Capelli, C., Hervig, T., Moreno-Estrada, A., Posukh, O. L., Balanovska, E., Balanovsky, O., Karachanak-Yankova, S., Sahakyan, H., Toncheva, D., Yepiskoposyan, L., Tyler-Smith, C., Xue, Y., Abdullah, M. S., Ruiz-Linares, A., Beall, C. M., Di Rienzo, A., Jeong, C., Starikovskaya, E. B., Metspalu, E., Parik, J., Villems, R., Henn, B. M., Hodoglugil, U., Mahley, R., Sajantila, A., Stamatoyannopoulos, G., Wee, J. T. S., Khusainova, R., Khusnutdinova, E., Litvinov, S., Ayodo, G., Comas, D., Hammer, M. F., Kivisild, T., Klitz, W., Winkler, C. A., Labuda, D., Bamshad, M., Jorde, L. B., Tishkoff, S. A., Watkins, W. S., Metspalu, M., Dryomov, S., Sukernik, R., Singh, L., Thangaraj, K., Pääbo, S., Kelso, J., Patterson, N., and Reich, D. (2016). The Simons Genome Diversity Project: 300 genomes from 142 diverse populations. Nature, 538(7624):201–206.

Marchi, N., Winkelbach, L., Schulz, I., Brami, M., Hofmanová, Z., Blöcher, J., Reyna-Blanco, C. S., Diekmann, Y., Thiéry, A., Kapopoulou, A., Link, V., Piuz, V., Kreutzer, S., Figarska, S. M., Ganiatsou, E., Pukaj, A., Struck, T. J., Gutenkunst, R. N., Karul, N., Gerritsen, F., Pechtl, J., Peters, J., Zeeb-Lanz, A., Lenneis, E., Teschler-Nicola, M., Triantaphyllou, S., Stefanović, S., Papageorgopoulou, C., Wegmann, D., Burger, J., and Excoffier, L. (2022). The genomic origins of the world’s first farmers. Cell, 185(11):1842–1859.e18.

Margaryan, A., Lawson, D. J., Sikora, M., Racimo, F., Rasmussen, S., Moltke, I., Cassidy, L. M., Jørsboe, E., Ingason, A., Pedersen, M. W., Korneliussen, T., Wilhelmson, H., Buś, M. M., De Barros Damgaard, P., Martiniano, R., Renaud, G., Bhérer, C., Moreno-Mayar, J. V., Fotakis, A. K., Allen, M., Allmäe, R., Molak, M., Cappellini, E., Scorrano, G., McColl, H., Buzhilova, A., Fox, A., Albrechtsen, A., Schütz, B., Skar, B., Arcini, C., Falys, C., Jonson, C. H., Błaszczyk, D., Pezhemsky, D., Turner-Walker, G., Gestsdóttir, H., Lundstrøm, I., Gustin, I., Mainland, I., Potekhina, I., Muntoni, I. M., Cheng, J., Stenderup, J., Ma, J., Gibson, J., Peets, J., Gustafsson, J., Iversen, K. H., Simpson, L., Strand, L., Loe, L., Sikora, M., Florek, M., Vretemark, M., Redknap, M., Bajka, M., Pushkina, T., Søvsø, M., Grigoreva, N., Christensen, T., Kastholm, O., Uldum, O., Favia, P., Holck, P., Sten, S., Arge, S. V., Ellingvåg, S., Moiseyev, V., Bogdanowicz, W., Magnusson, Y., Orlando, L., Pentz, P., Jessen, M. D., Pedersen, A., Collard, M., Bradley, D. G., Jørkov, M. L., Arneborg, J., Lynnerup, N., Price, N., Gilbert, M. T. P., Allentoft, M. E., Bill, J., Sindbæk, S. M., Hedeager, L., Kristiansen, K., Nielsen, R., Werge, T., and Willerslev, E. (2020). Population genomics of the Viking world. Nature, 585(7825):390–396.

Mathieson, I., Alpaslan-Roodenberg, S., Posth, C., Szécsényi-Nagy, A., Rohland, N., Mallick, S., Olalde, I., Broomandkhoshbacht, N., Candilio, F., Cheronet, O., Fernandes, D., Ferry, M., Gamarra, B., Fortes, G. G., Haak, W., Harney, E., Jones, E., Keating, D., Krause-Kyora, B., Kucukkalipci, I., Michel, M., Mittnik, A., Nägele, K., Novak, M., Oppenheimer, J., Patterson, N., Pfrengle, S., Sirak, K., Stewardson, K., Vai, S., Alexandrov, S., Alt, K. W., Andreescu, R., Antonović, D., Ash, A., Atanassova, N., Bacvarov, K., Gusztáv, M. B., Bocherens, H., Bolus, M., Boroneanţ, A., Boyadzhiev, Y., Budnik, A., Burmaz, J., Chohadzhiev, S., Conard, N. J., Cottiaux, R., Čuka, M., Cupillard, C., Drucker, D. G., Elenski, N., Francken, M., Galabova, B., Ganetsovski, G., Gély, B., Hajdu, T., Handzhyiska, V., Harvati, K., Higham, T., Iliev, S., Janković, I., Karavanić, I., Kennett, D. J., Komšo, D., Kozak, A., Labuda, D., Lari, M., Lazar, C., Leppek, M., Leshtakov, K., Vetro, D. L., Los, D., Lozanov, I., Malina, M., Martini, F., McSweeney, K., Meller, H., Menđušić, M., Mirea, P., Moiseyev, V., Petrova, V., Price, T. D., Simalcsik, A., Sineo, L., Šlaus, M., Slavchev, V., Stanev, P., Starović, A., Szeniczey, T., Talamo, S., Teschler-Nicola, M., Thevenet, C., Valchev, I., Valentin, F., Vasilyev, S., Veljanovska, F., Venelinova, S., Veselovskaya, E., Viola, B., Virag, C., Zaninović, J., Zäuner, S., Stockhammer, P. W., Catalano, G., Krauß, R., Caramelli, D., Zariņa, G., Gaydarska, B., Lillie, M., Nikitin, A. G., Potekhina, I., Papathanasiou, A., Borić, D., Bonsall, C., Krause, J., Pinhasi, R., and Reich, D. (2018). The genomic history of southeastern Europe. Nature, 555(7695):197–203.

Mathieson, I., Lazaridis, I., Rohland, N., Mallick, S., Patterson, N., Roodenberg, S. A., Harney, E., Stewardson, K., Fernandes, D., Novak, M., Sirak, K., Gamba, C., Jones, E. R., Llamas, B., Dryomov, S., Pickrell, J., Arsuaga, J. L., De Castro, J. M. B., Carbonell, E., Gerritsen, F., Khokhlov, A., Kuznetsov, P., Lozano, M., Meller, H., Mochalov, O., Moiseyev, V., Guerra, M. A. R., Roodenberg, J., Vergès, J. M., Krause, J., Cooper, A., Alt, K. W., Brown, D., Anthony, D., Lalueza-Fox, C., Haak, W., Pinhasi, R., and Reich, D. (2015). Genome-wide patterns of selection in 230 ancient Eurasians. Nature, 528(7583):499–503.

Mittnik, A., Wang, C.-C., Svoboda, J., and Krause, J. (2016). A Molecular Approach to the Sexing of the Triple Burial at the Upper Paleolithic Site of Dolní Věstonice. PLOS ONE, 11(10):e0163019.

Narasimhan, V. M., Patterson, N., Moorjani, P., Rohland, N., Bernardos, R., Mallick, S., Lazaridis, I., Nakatsuka, N., Olalde, I., Lipson, M., Kim, A. M., Olivieri, L. M., Coppa, A., Vidale, M., Mallory, J., Moiseyev, V., Kitov, E., Monge, J., Adamski, N., Alex, N., Broomandkhoshbacht, N., Candilio, F., Callan, K., Cheronet, O., Culleton, B. J., Ferry, M., Fernandes, D., Freilich, S., Gamarra, B., Gaudio, D., Hajdinjak, M., Harney, É., Harper, T. K., Keating, D., Lawson, A. M., Mah, M., Mandl, K., Michel, M., Novak, M., Oppenheimer, J., Rai, N., Sirak, K., Slon, V., Stewardson, K., Zalzala, F., Zhang, Z., Akhatov, G., Bagashev, A. N., Bagnera, A., Baitanayev, B., Bendezu-Sarmiento, J., Bissembaev, A. A., Bonora, G. L., Chargynov, T. T., Chikisheva, T., Dashkovskiy, P. K., Derevianko, A., Dobeš, M., Douka, K., Dubova, N., Duisengali, M. N., Enshin, D., Epimakhov, A., Fribus, A. V., Fuller, D., Goryachev, A., Gromov, A., Grushin, S. P., Hanks, B., Judd, M., Kazizov, E., Khokhlov, A., Krygin, A. P., Kupriyanova, E., Kuznetsov, P., Luiselli, D., Maksudov, F., Mamedov, A. M., Mamirov, T. B., Meiklejohn, C., Merrett, D. C., Micheli, R., Mochalov, O., Mustafokulov, S., Nayak, A., Pettener, D., Potts, R., Razhev, D., Rykun, M., Sarno, S., Savenkova, T. M., Sikhymbaeva, K., Slepchenko, S. M., Soltobaev, O. A., Stepanova, N., Svyatko, S., Tabaldiev, K., Teschler-Nicola, M., Tishkin, A. A., Tkachev, V. V., Vasilyev, S., Velemínský, P., Voyakin, D., Yermolayeva, A., Zahir, M., Zubkov, V. S., Zubova, A., Shinde, V. S., Lalueza-Fox, C., Meyer, M., Anthony, D., Boivin, N., Thangaraj, K., Kennett, D. J., Frachetti, M., Pinhasi, R., and Reich, D. (2019). The formation of human populations in South and Central Asia. Science, 365(6457):eaat7487.

Nassar, L. R., Barber, G. P., Benet-Pagès, A., Casper, J., Clawson, H., Diekhans, M., Fischer, C., Gonzalez, J. N., Hinrichs, A. S., Lee, B. T., Lee, C. M., Muthuraman, P., Nguy, B., Pereira, T., Nejad, P., Perez, G., Raney, B. J., Schmelter, D., Speir, M. L., Wick, B. D., Zweig, A. S., Haussler, D., Kuhn, R. M., Haeussler, M., and Kent, W. J. (2023). The UCSC Genome Browser database: 2023 update. Nucleic Acids Research, 51(D1):D1188–D1195.

Novak, M., Olalde, I., Ringbauer, H., Rohland, N., Ahern, J., Balen, J., Janković, I., Potrebica, H., Pinhasi, R., and Reich, D. (2021). Genome-wide analysis of nearly all the victims of a 6200 year old massacre. PLOS ONE, 16(3):e0247332.

Nursyifa, C., Brüniche-Olsen, A., Garcia-Erill, G., Heller, R., and Albrechtsen, A. (2022). Joint identification of sex and sex-linked scaffolds in non-model organisms using low depth sequencing data. Molecular Ecology Resources, 22(2):458–467.

Olalde, I., Brace, S., Allentoft, M. E., Armit, I., Kristiansen, K., Booth, T., Rohland, N., Mallick, S., Szécsényi-Nagy, A., Mittnik, A., Altena, E., Lipson, M., Lazaridis, I., Harper, T. K., Patterson, N., Broomandkhoshbacht, N., Diekmann, Y., Faltyskova, Z., Fernandes, D., Ferry, M., Harney, E., De Knijff, P., Michel, M., Oppenheimer, J., Stewardson, K., Barclay, A., Alt, K. W., Liesau, C., Ríos, P., Blasco, C., Miguel, J. V., García, R. M., Fernández, A. A., Bánffy, E., Bernabò-Brea, M., Billoin, D., Bonsall, C., Bonsall, L., Allen, T., Büster, L., Carver, S., Navarro, L. C., Craig, O. E., Cook, G. T., Cunliffe, B., Denaire, A., Dinwiddy, K. E., Dodwell, N., Ernée, M., Evans, C., Kuchařík, M., Farré, J. F., Fowler, C., Gazenbeek, M., Pena, R. G., Haber-Uriarte, M., Haduch, E., Hey, G., Jowett, N., Knowles, T., Massy, K., Pfrengle, S., Lefranc, P., Lemercier, O., Lefebvre, A., Martínez, C. H., Olmo, V. G., Ramírez, A. B., Maurandi, J. L., Majó, T., McKinley, J. I., McSweeney, K., Mende, B. G., Modi, A., Kulcsár, G., Kiss, V., Czene, A., Patay, R., Endrődi, A., Köhler, K., Hajdu, T., Szeniczey, T., Dani, J., Bernert, Z., Hoole, M., Cheronet, O., Keating, D., Velemínský, P., Dobeš, M., Candilio, F., Brown, F., Fernández, R. F., Herrero-Corral, A.-M., Tusa, S., Carnieri, E., Lentini, L., Valenti, A., Zanini, A., Waddington, C., Delibes, G., Guerra-Doce, E., Neil, B., Brittain, M., Luke, M., Mortimer, R., Desideri, J., Besse, M., Brücken, G., Furmanek, M., Hałuszko, A., Mackiewicz, M., Rapiński, A., Leach, S., Soriano, I., Lillios, K. T., Cardoso, J. L., Pearson, M. P., Włodarczak, P., Price, T. D., Prieto, P., Rey, P.-J., Risch, R., Rojo Guerra, M. A., Schmitt, A., Serralongue, J., Silva, A. M., Smrčka, V., Vergnaud, L., Zilhão, J., Caramelli, D., Higham, T., Thomas, M. G., Kennett, D. J., Fokkens, H., Heyd, V., Sheridan, A., Sjögren, K.-G., Stockhammer, P. W., Krause, J., Pinhasi, R., Haak, W., Barnes, I., Lalueza-Fox, C., and Reich, D. (2018). The Beaker phenomenon and the genomic transformation of northwest Europe. Nature, 555(7695):190–196.

Olalde, I., Mallick, S., Patterson, N., Rohland, N., Villalba-Mouco, V., Silva, M., Dulias, K., Edwards, C. J., Gandini, F., Pala, M., Soares, P., Ferrando-Bernal, M., Adamski, N., Broomandkhoshbacht, N., Cheronet, O., Culleton, B. J., Fernandes, D., Lawson, A. M., Mah, M., Oppenheimer, J., Stewardson, K., Zhang, Z., Jiménez Arenas, J. M., Toro Moyano, I. J., Salazar-García, D. C., Castanyer, P., Santos, M., Tremoleda, J., Lozano, M., García Borja, P., Fernández-Eraso, J., Mujika-Alustiza, J. A., Barroso, C., Bermúdez, F. J., Viguera Mínguez, E., Burch, J., Coromina, N., Vivó, D., Cebrià, A., Fullola, J. M., García-Puchol, O., Morales, J. I., Oms, F. X., Majó, T., Vergès, J. M., Díaz-Carvajal, A., Ollich-Castanyer, I., López-Cachero, F. J., Silva, A. M., Alonso-Fernández, C., Delibes De Castro, G., Jiménez Echevarría, J., Moreno-Márquez, A., Pascual Berlanga, G., Ramos-García, P., Ramos-Muñoz, J., Vijande Vila, E., Aguilella Arzo, G., Esparza Arroyo, Á., Lillios, K. T., Mack, J., Velasco-Vázquez, J., Waterman, A., Benítez De Lugo Enrich, L., Benito Sánchez, M., Agustí, B., Codina, F., De Prado, G., Estalrrich, A., Fernández Flores, Á., Finlayson, C., Finlayson, G., Finlayson, S., Giles-Guzmán, F., Rosas, A., Barciela González, V., García Atiénzar, G., Hernández Pérez, M. S., Llanos, A., Carrión Marco, Y., Collado Beneyto, I., López-Serrano, D., Sanz Tormo, M., Valera, A. C., Blasco, C., Liesau, C., Ríos, P., Daura, J., De Pedro Michó, M. J., Diez-Castillo, A. A., Flores Fernández, R., Francès Farré, J., Garrido-Pena, R., Gonçalves, V. S., Guerra-Doce, E., Herrero-Corral, A. M., Juan-Cabanilles, J., López-Reyes, D., McClure, S. B., Merino Pérez, M., Oliver Foix, A., Sanz Borràs, M., Sousa, A. C., Vidal Encinas, J. M., Kennett, D. J., Richards, M. B., Werner Alt, K., Haak, W., Pinhasi, R., Lalueza-Fox, C., and Reich, D. (2019). The genomic history of the Iberian Peninsula over the past 8000 years. Science, 363(6432):1230–1234.

Palmer, D. H., Rogers, T. F., Dean, R., and Wright, A. E. (2019). How to identify sex chromosomes and their turnover. Molecular Ecology, 28(21):4709–4724.

Patterson, N., Isakov, M., Booth, T., Büster, L., Fischer, C.-E., Olalde, I., Ringbauer, H., Akbari, A., Cheronet, O., Bleasdale, M., Adamski, N., Altena, E., Bernardos, R., Brace, S., Broomandkhoshbacht, N., Callan, K., Candilio, F., Culleton, B., Curtis, E., Demetz, L., Carlson, K. S. D., Edwards, C. J., Fernandes, D. M., Foody, M. G. B., Freilich, S., Goodchild, H., Kearns, A., Lawson, A. M., Lazaridis, I., Mah, M., Mallick, S., Mandl, K., Micco, A., Michel, M., Morante, G. B., Oppenheimer, J., Özdoğan, K. T., Qiu, L., Schattke, C., Stewardson, K., Workman, J. N., Zalzala, F., Zhang, Z., Agustí, B., Allen, T., Almássy, K., Amkreutz, L., Ash, A., Baillif-Ducros, C., Barclay, A., Bartosiewicz, L., Baxter, K., Bernert, Z., Blažek, J., Bodružić, M., Boissinot, P., Bonsall, C., Bradley, P., Brittain, M., Brookes, A., Brown, F., Brown, L., Brunning, R., Budd, C., Burmaz, J., Canet, S., Carnicero-Cáceres, S., Čaušević-Bully, M., Chamberlain, A., Chauvin, S., Clough, S., Čondić, N., Coppa, A., Craig, O., Črešnar, M., Cummings, V., Czifra, S., Danielisová, A., Daniels, R., Davies, A., De Jersey, P., Deacon, J., Deminger, C., Ditchfield, P. W., Dizdar, M., Dobeš, M., Dobisíková, M., Domboróczki, L., Drinkall, G., Ðukić, A., Ernée, M., Evans, C., Evans, J., Fernández-Götz, M., Filipović, S., Fitzpatrick, A., Fokkens, H., Fowler, C., Fox, A., Gallina, Z., Gamble, M., González Morales, M. R., González-Rabanal, B., Green, A., Gyenesei, K., Habermehl, D., Hajdu, T., Hamilton, D., Harris, J., Hayden, C., Hendriks, J., Hernu, B., Hey, G., Horňák, M., Ilon, G., Istvánovits, E., Jones, A. M., Kavur, M. B., Kazek, K., Kenyon, R. A., Khreisheh, A., Kiss, V., Kleijne, J., Knight, M., Kootker, L. M., Kovács, P. F., Kozubová, A., Kulcsár, G., Kulcsár, V., Le Pennec, C., Legge, M., Leivers, M., Loe, L., López-Costas, O., Lord, T., Los, D., Lyall, J., Marín-Arroyo, A. B., Mason, P., Matošević, D., Maxted, A., McIntyre, L., McKinley, J., McSweeney, K., Meijlink, B., Mende, B. G., Menđušić, M., Metlička, M., Meyer, S., Mihovilić, K., Milasinovic, L., Minnitt, S., Moore, J., Morley, G., Mullan, G., Musilová, M., Neil, B., Nicholls, R., Novak, M., Pala, M., Papworth, M., Paresys, C., Patten, R., Perkić, D., Pesti, K., Petit, A., Petriščáková, K., Pichon, C., Pickard, C., Pilling, Z., Price, T. D., Radović, S., Redfern, R., Resutík, B., Rhodes, D. T., Richards, M. B., Roberts, A., Roefstra, J., Sankot, P., Šefčáková, A., Sheridan, A., Skae, S., Šmolíková, M., Somogyi, K., Somogyvári, Á., Stephens, M., Szabó, G., Szécsényi-Nagy, A., Szeniczey, T., Tabor, J., Tankó, K., Maria, C. T., Terry, R., Teržan, B., Teschler-Nicola, M., Torres-Martínez, J. F., Trapp, J., Turle, R., Ujvári, F., Van Der Heiden, M., Veleminsky, P., Veselka, B., Vytlačil, Z., Waddington, C., Ware, P., Wilkinson, P., Wilson, L., Wiseman, R., Young, E., Zaninović, J., Žitňan, A., Lalueza-Fox, C., De Knijff, P., Barnes, I., Halkon, P., Thomas, M. G., Kennett, D. J., Cunliffe, B., Lillie, M., Rohland, N., Pinhasi, R., Armit, I., and Reich, D. (2022). Large-scale migration into Britain during the Middle to Late Bronze Age. Nature, 601(7894):588–594.

Pečnerová, P., Díez-del-Molino, D., Dussex, N., Feuerborn, T., Von Seth, J., Van Der Plicht, J., Nikolskiy, P., Tikhonov, A., Vartanyan, S., and Dalén, L. (2017). Genome-Based Sexing Provides Clues about Behavior and Social Structure in the Woolly Mammoth. Current Biology, 27(22):3505–3510.e3.

Peppin, L., McEwing, R., Ogden, R., Hermes, R., Harper, C., Guthrie, A., and Carvalho, G. R. (2010). Molecular sexing of African rhinoceros. Conservation Genetics, 11(3):1181–1184.

Prendergast, M. E., Lipson, M., Sawchuk, E. A., Olalde, I., Ogola, C. A., Rohland, N., Sirak, K. A., Adamski, N., Bernardos, R., Broomandkhoshbacht, N., Callan, K., Culleton, B. J., Eccles, L., Harper, T. K., Lawson, A. M., Mah, M., Oppenheimer, J., Stewardson, K., Zalzala, F., Ambrose, S. H., Ayodo, G., Gates, H. L., Gidna, A. O., Katongo, M., Kwekason, A., Mabulla, A. Z. P., Mudenda, G. S., Ndiema, E. K., Nelson, C., Robertshaw, P., Kennett, D. J., Manthi, F. K., and Reich, D. (2019). Ancient DNA reveals a multistep spread of the first herders into sub-Saharan Africa. Science, 365(6448):eaaw6275.

Rangavittal, S., Stopa, N., Tomaszkiewicz, M., Sahlin, K., Makova, K. D., and Medvedev, P. (2019). DiscoverY: A classifier for identifying Y chromosome sequences in male assemblies. BMC Genomics, 20(1):641.

Reitsema, L. J., Mittnik, A., Kyle, B., Catalano, G., Fabbri, P. F., Kazmi, A. C. S., Reinberger, K. L., Sineo, L., Vassallo, S., Bernardos, R., Broomandkhoshbacht, N., Callan, K., Candilio, F., Cheronet, O., Curtis, E., Fernandes, D., Lari, M., Lawson, A. M., Mah, M., Mallick, S., Mandl, K., Micco, A., Modi, A., Oppenheimer, J., Özdogan, K. T., Rohland, N., Stewardson, K., Vai, S., Vergata, C., Workman, J. N., Zalzala, F., Zaro, V., Achilli, A., Anagnostopoulos, A., Capelli, C., Constantinou, V., Lancioni, H., Olivieri, A., Papadopoulou, A., Psatha, N., Semino, O., Stamatoyannopoulos, J., Valliannou, I., Yannaki, E., Lazaridis, I., Patterson, N., Ringbauer, H., Caramelli, D., Pinhasi, R., and Reich, D. (2022). The diverse genetic origins of a Classical period Greek army. Proceedings of the National Academy of Sciences, 119(41):e2205272119.

Rivollat, M., Jeong, C., Schiffels, S., Küçükkalıpçı, İ., Pemonge, M.-H., Rohrlach, A. B., Alt, K. W., Binder, D., Friederich, S., Ghesquière, E., Gronenborn, D., Laporte, L., Lefranc, P., Meller, H., Réveillas, H., Rosenstock, E., Rottier, S., Scarre, C., Soler, L., Wahl, J., Krause, J., Deguilloux, M.-F., and Haak, W. (2020). Ancient genome-wide DNA from France highlights the complexity of interactions between Mesolithic hunter-gatherers and Neolithic farmers. Science Advances, 6(22):eaaz5344.

Robledo-Ruiz, D. A., Austin, L., Amos, J. N., Castrejón-Figueroa, J., Harley, D. K. P., Magrath, M. J. L., Sunnucks, P., and Pavlova, A. (2023). Easy-to-use R functions to separate reduced-representation genomic datasets into sex-linked and autosomal loci, and conduct sex assignment. Molecular Ecology Resources, pages 1755–0998.13844.

Sastre, N., Francino, O., Lampreave, G., Bologov, V. V., López-Martín, J. M., Sánchez, A., and Ramírez, O. (2009). Sex identification of wolf (Canis lupus) using non-invasive samples. Conservation Genetics, 10(3):555–558.

Scheib, C. L., Li, H., Desai, T., Link, V., Kendall, C., Dewar, G., Griffith, P. W., Mörseburg, A., Johnson, J. R., Potter, A., Kerr, S. L., Endicott, P., Lindo, J., Haber, M., Xue, Y., Tyler-Smith, C., Sandhu, M. S., Lorenz, J. G., Randall, T. D., Faltyskova, Z., Pagani, L., Danecek, P., O’Connell, T. C., Martz, P., Boraas, A. S., Byrd, B. F., Leventhal, A., Cambra, R., Williamson, R., Lesage, L., Holguin, B., Ygnacio-De Soto, E., Rosas, J., Metspalu, M., Stock, J. T., Manica, A., Scally, A., Wegmann, D., Malhi, R. S., and Kivisild, T. (2018). Ancient human parallel lineages within North America contributed to a coastal expansion. Science, 360(6392):1024–1027.

Schurz, H., Salie, M., Tromp, G., Hoal, E. G., Kinnear, C. J., and Möller, M. (2019). The X chromosome and sex-specific effects in infectious disease susceptibility. Human Genomics, 13(1):2.

Sigeman, H., Sinclair, B., and Hansson, B. (2022). Findzx: An automated pipeline for detecting and visualising sex chromosomes using whole-genome sequencing data. BMC Genomics, 23(1):328.

Sirak, K. A., Fernandes, D. M., Lipson, M., Mallick, S., Mah, M., Olalde, I., Ringbauer, H., Rohland, N., Hadden, C. S., Harney, É., Adamski, N., Bernardos, R., Broomandkhoshbacht, N., Callan, K., Ferry, M., Lawson, A. M., Michel, M., Oppenheimer, J., Stewardson, K., Zalzala, F., Patterson, N., Pinhasi, R., Thompson, J. C., Van Gerven, D., and Reich, D. (2021). Social stratification without genetic differentiation at the site of Kulubnarti in Christian Period Nubia. Nature Communications, 12(1):7283.

Skoglund, P., Storå, J., Götherström, A., and Jakobsson, M. (2013). Accurate sex identification of ancient human remains using DNA shotgun sequencing. Journal of Archaeological Science, 40(12):4477–4482.

Skuse, D., Printzlau, F., and Wolstencroft, J. (2018). Chapter 24 - sex chromosome aneuploidies. In Geschwind, D. H., Paulson, H. L., and Klein, C., editors, Neurogenetics, Part I, volume 147 of Handbook of Clinical Neurology, pages 355–376. Elsevier.

Stöck, M., Kratochvíl, L., Kuhl, H., Rovatsos, M., Evans, B. J., Suh, A., Valenzuela, N., Veyrunes, F., Zhou, Q., Gamble, T., Capel, B., Schartl, M., and Guiguen, Y. (2021). A brief review of vertebrate sex evolution with a pledge for integrative research: Towards ‘ sexomics ‘. Philosophical Transactions of the Royal Society B: Biological Sciences, 376(1832):20200426.

The Tree of Sex Consortium, Ashman, T.-L., Bachtrog, D., Blackmon, H., Goldberg, E. E., Hahn, M. W., Kirkpatrick, M., Kitano, J., Mank, J. E., Mayrose, I., Ming, R., Otto, S. P., Peichel, C. L., Pennell, M. W., Perrin, N., Ross, L., Valenzuela, N., and Vamosi, J. C. (2014). Tree of Sex: A database of sexual systems. Scientific Data, 1(1):140015.

Tiesler, V., Sedig, J., Nakatsuka, N., Mallick, S., Lazaridis, I., Bernardos, R., Broomandkhoshbacht, N., Oppenheimer, J., Lawson, A. M., Stewardson, K., Rohland, N., Kennett, D. J., Price, T. D., and Reich, D. (2022). Life and death in early colonial Campeche: New insights from ancient DNA. Antiquity, 96(388):937–954.

Veeramah, K. R., Rott, A., Groß, M., Van Dorp, L., López, S., Kirsanow, K., Sell, C., Blöcher, J., Wegmann, D., Link, V., Hofmanová, Z., Peters, J., Trautmann, B., Gairhos, A., Haberstroh, J., Päffgen, B., Hellenthal, G., Haas-Gebhard, B., Harbeck, M., and Burger, J. (2018). Population genomic analysis of elongated skulls reveals extensive female-biased immigration in Early Medieval Bavaria. Proceedings of the National Academy of Sciences, 115(13):3494–3499.

Villalba-Mouco, V., Oliart, C., Rihuete-Herrada, C., Childebayeva, A., Rohrlach, A. B., Fregeiro, M. I., Celdrán Beltrán, E., Velasco-Felipe, C., Aron, F., Himmel, M., Freund, C., Alt, K. W., Salazar-García, D. C., García Atiénzar, G., De Miguel Ibáñez, M. P., Hernández Pérez, M. S., Barciela, V., Romero, A., Ponce, J., Martínez, A., Lomba, J., Soler, J., Martínez, A. P., Avilés Fernández, A., Haber-Uriarte, M., Roca De Togores Muñoz, C., Olalde, I., Lalueza-Fox, C., Reich, D., Krause, J., García Sanjuán, L., Lull, V., Micó, R., Risch, R., and Haak, W. (2021). Genomic transformation and social organization during the Copper Age–Bronze Age transition in southern Iberia. Science Advances, 7(47):eabi7038.

Žegarac, A., Winkelbach, L., Blöcher, J., Diekmann, Y., Krečković Gavrilović, M., Porčić, M., Stojković, B., Milašinović, L., Schreiber, M., Wegmann, D., Veeramah, K. R., Stefanović, S., and Burger, J. (2021). Ancient genomes provide insights into family structure and the heredity of social status in the early Bronze Age of southeastern Europe. Scientific Reports, 11(1):10072.

