## Supplementary Information for "Accurate Bayesian inference of sex chromosome karyotypes and sex-linked scaffolds from low-depth sequencing data"

#### 1 Model

We aim at jointly inferring the genomic sex $\boldsymbol{s} = (s_1, \ldots, s_I)$, the ploidy-related parameters $\boldsymbol{\rho} = (\rho_1, \ldots, \rho_C)$, the $M$ vectors of mapping attractors $(\boldsymbol{\gamma}_1, \ldots, \boldsymbol{\gamma}_M)$, the variances $\boldsymbol{\sigma} = (\sigma_1, \ldots, \sigma_M)$, the autosomal trisomy indicators $\boldsymbol{z} = (z_{11}, \ldots, z_{1C}, z_{21}, \ldots, z_{IC})$ as well as the error terms $\boldsymbol{\epsilon} = (\epsilon_1, \ldots, \epsilon_R)$ using a Bayesian scheme. Figure 1 shows the directed acyclic graph (DAG) of the model that visualizes the assumptions about independence and conditional independence.

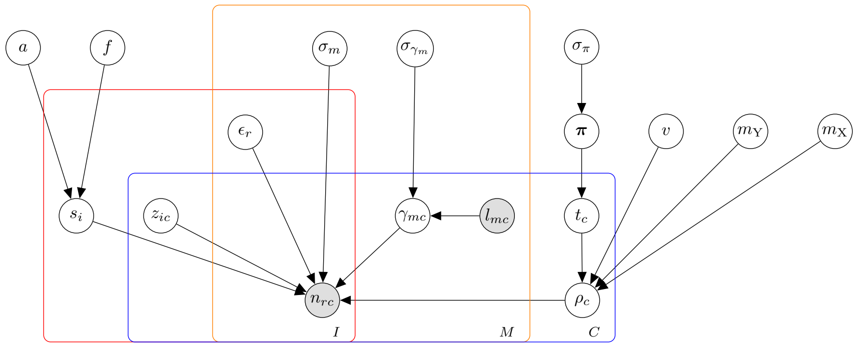


Figure 1: Directed acyclic graph (DAG) of the model. The shaded nodes represent observations, i.e. the read counts $n_{rc}$ and the reference statistics $l_{mc}$. Only prior parameters that are inferred are drawn. Recall that the index $r = (i, m)$ indicates the combination of indices $i$ and $m$.

#### 2 Prior distributions

**Prior on $s_i$** This prior is described in the main text. The relative frequencies of aneuploid karyotypes were calculated from Skuse et al. (2018) as $\boldsymbol{v} = (v_{\mathrm{X0}}, \ldots, v_{\mathrm{XXYY}}) = (0.086, 0.432, 0.216, 0.254, 0.012)$.

**Prior on $a$ and $f$** We assume that $a$ and $f$ both follow a Beta distribution, $a \sim \mathrm{Beta}(\alpha_a, \beta_a)$ and $f \sim \mathrm{Beta}(\alpha_f, \beta_f)$. We re-parametrize the Beta distributions in terms of mean and variance and use 1e-4, 1e-10 for $a$ and 0.5, 0.02 for $f$ as default values for mean and variance of the Beta distributions, respectively.

**Prior on $z_{ic}$** We assume that $z_{ic}$ follow an Bernouilli distribution $z_{ic} \sim \mathrm{Bernouilli}(\pi_z)$ with a default $\pi_z = \frac{1}{1000}$.

**Prior on $\epsilon_r$** We assume that $\epsilon_r$ follows an exponential prior $\epsilon_r \sim \mathrm{Exp}(\lambda_\epsilon)$ with a default $\lambda_\epsilon = 1000$.

**Prior on $\sigma_m$** Since the variance of the Dirichlet distribution $\sigma_m$ can be very small, we infer the logarithm of $\sigma_m$ and assume $\log \sigma_m \sim \mathcal{U}(-\infty, \infty)$.

**Prior on $\boldsymbol{\gamma}_m$** We assume that each vector $\boldsymbol{\gamma}_m = (\gamma_{m1}, \ldots, \gamma_{mC})$ follows a Dirichlet distribution $\boldsymbol{\gamma}_m \sim \text{Dir}(\sigma_{\gamma_m}^{-1}\boldsymbol{l}_m)$, where $\boldsymbol{l}_m$ is the vector of expected read proportions for each scaffold, $\boldsymbol{l}_m = (l_1, \ldots, l_C)$, and $\sigma_{\gamma_m}$ controls the variance of the Dirichlet distribution. The expected read proportions are given by $l_{mc} = \frac{L_{mc}}{\sum_g L_{mg}}$, where $L_{mc}$ denote the expected number of reads mapping to scaffold $c$ for method $m$. The expected number of reads mapping to each scaffold depends on the sequencing method, and it is up to the user to define meaningful expectations. For instance, we recommend to use the length of the scaffold in the case of whole-genome shotgun sequencing and the number of markers per scaffold in the case of target enrichment capture.

**Prior on $\sigma_{\gamma_m}$** We assume that $\sigma_{\gamma_m}$ follows an exponential prior $\sigma_{\gamma_m} \sim \text{Exp}(\lambda_{\sigma_{\gamma_m}})$ with a default $\lambda_{\sigma_{\gamma_m}} = 1000$.

**Prior on $\rho_c$** We assume a mixture of four Beta distributions as a prior on $\rho_c$. The scaffold type $t_c = \text{A}, \text{Y}, \text{X}, \text{D}$ indicates the mixture component for scaffold $c$.

For the autosomes ($t_c = \text{A}$), we use a symmetrical Beta distribution $\rho_c | t_c = \text{A} \sim \text{Beta}(\mu, \mu)$ with $\mu > 1$, which has an expected value $\mathbb{E}(\rho_c) = \frac{1}{2}$. Our goal is to define a prior distribution on $\mu$ that pushes towards a "peaky" Beta distribution. To achieve this, we re-parametrize the symmetric Beta distribution and infer its variance $v$, which is related to $\mu$ as $\mu = \frac{1}{8v} - \frac{1}{2}$. In order to make sure that $\mu > 1$, we impose the constraint that $v < \frac{1}{12}$. We assume that $v$ follows an exponential distribution, $v \sim \text{Exp}(\lambda_v)$ and use and use $\lambda_v = 10000$ as a default value.

For the Y-linked scaffolds ($t_c = \text{Y}$), we use a left-leaning distribution $\rho_c | t_c = \text{Y} \sim \text{Beta}(1, \beta)$ with $\beta > 1$. Our goal is to define a prior distribution on $\beta$ that pushes the Beta distribution towards zero. To achieve this, we re-parametrize the Beta distribution and the infer its mean $m_\text{Y}$, which is related to $\beta$ as $\beta = \frac{1}{m_\text{Y}} - 1$ if $\alpha = 1$. In order to make sure that $\beta > 1$, we impose the constraint that $m_\text{Y} < \frac{1}{2}$. We assume that $m_\text{Y}$ follows an exponential distribution, $m_\text{Y} \sim \text{Exp}(\lambda_{m_\text{Y}})$ and use $\lambda_{m_\text{Y}} = 1000$ as a default value.

For the X-linked scaffolds ($t_c = \text{X}$), we use a right-leaning distribution $\rho_c | t_c = \text{X} \sim \text{Beta}(\alpha, 1)$ with $\alpha > 1$. Our goal is to define a prior distribution on $\alpha$ that pushes the Beta distribution towards one. To achieve this, we re-parametrize the Beta distribution and the infer a parameter $m_\text{X}$, which corresponds to the flipped mean $= 1 - m_\text{X}$ and which is related to $\alpha$ as $\alpha = \frac{1}{1 - m_\text{X}} - 1$ if $\beta = 1$. In order to make sure that $\alpha > 1$, we impose the constraint that $m_\text{X} < \frac{1}{2}$. We assume that $m_\text{X}$ follows an exponential distribution, $m_\text{X} \sim \text{Exp}(\lambda_{m_\text{X}})$ and use $\lambda_{m_\text{X}} = 10000$ as a default value.

Finally, the non-canonical scaffolds of type $t_c = \text{D}$ are modelled using a flat Beta distribution $\rho_c | t_c = \text{D} \sim \text{Beta}(1, 1)$.

**Prior on $t_c$** We assume that scaffold types $t_c$ follow a Categorical distribution $t_c \sim \text{Cat}(\boldsymbol{\pi})$ where $\boldsymbol{\pi}$ is a vector of probabilities for each category, $\boldsymbol{\pi} = (\pi_\text{A}, \pi_\text{Y}, \pi_\text{X}, \pi_\text{D})$.

**Prior on $\boldsymbol{\pi}$** We assume that the probability vector $\boldsymbol{\pi} = (\pi_\text{A}, \pi_\text{Y}, \pi_\text{X}, \pi_\text{D})$ follows a Dirichlet prior as $\boldsymbol{\pi} \sim \text{Dir}(\tilde{\alpha}_\text{A}, \tilde{\alpha}_\text{Y}, \tilde{\alpha}_\text{X}, \tilde{\alpha}_\text{D})$. Here, each parameter $\tilde{\alpha}_T$ for $T = \{\text{A}, \text{Y}, \text{X}, \text{D}\}$ is given by $\tilde{\alpha}_T = \alpha_T \sigma_\pi$, where $\alpha_T$ is a fixed value that represents the expected value of $\pi_T$, and $\sigma_\pi$ is an inferred parameter that models the variance of the Dirichlet distribution. We use $\alpha_\text{A} = 44, \alpha_\text{Y} = 1, \alpha_\text{X} = 1$ and $\alpha_\text{D} = 0.1$ by default.

**Prior on $\sigma_\pi$** We assume that $\sigma_\pi$ follows an exponential distribution $\sigma_\pi \sim \text{Exp}(\lambda_{\sigma_\pi})$ with a default $\lambda_{\sigma_\pi} = 10$.

### 3 Bayesian inference

We use an Markov chain Monte Carlo (MCMC) scheme to generate samples from the posterior $\mathbb{P}(\boldsymbol{s}, \boldsymbol{\epsilon}, \boldsymbol{\sigma}, \boldsymbol{\gamma}, \boldsymbol{\rho}, a, f, \sigma_\gamma, \boldsymbol{t}, \boldsymbol{\pi}, \sigma_\pi, v, m_\text{Y}, m_\text{X} | \boldsymbol{N})$, where $\boldsymbol{N}$ denotes the $R \times C$ matrix of read counts with entries $[\boldsymbol{N}]_{rc} = n_{rc}$.

The likelihood of a vector $\boldsymbol{n}_r = (n_{r1}, \ldots, n_{rC})$ is the density of the Dirichlet-Multinomial distribution:

$$\mathbb{P}(\boldsymbol{n}_r|N_r, \boldsymbol{\alpha}(r)) = \frac{\Gamma\left(\sum_c \alpha_c(r)\right)\Gamma\left(N_r+1\right)}{\Gamma\left(N_r+\sum_c \alpha_c(r)\right)} \prod_{c=1}^{C} \frac{\Gamma\left(n_{rc}+\alpha_c(r)\right)}{\Gamma\left(\alpha_c(r)\right)\Gamma\left(n_{rc}+1\right)},$$

where $\Gamma$ denotes the Gamma function. Considering $n_{rc}$ and $N_r = \sum_c n_{rc}$ as constant, the terms $\Gamma\left(N_r+1\right)$ and $\Gamma\left(n_{rc}+1\right)$ are normalization constants and can be ignored when evaluating Hastings ratios. We thus get the following relevant log likelihood:

$$\log \mathbb{P}(\boldsymbol{n}_r|N_r, \boldsymbol{\alpha}(r)) = \ln\Gamma\left(\sum_{c=1}^{C}\alpha_c(r)\right) - \sum_{c=1}^{C}\ln\Gamma\left(\alpha_c(r)\right) + \sum_{c=1}^{C}\ln\Gamma\left(n_{rc}+\alpha_c(r)\right) - \ln\Gamma\left(N_r + \sum_{c=1}^{C}\alpha_c(r)\right).$$

The log Hastings ratio $\log h_\theta$ for an update of a parameter $\theta = \{s_i, \epsilon_r, \sigma_m, \gamma_{mc}, \rho_c\}$ is given by

$$\log h_\theta = \log \mathbb{P}(\theta') - \log \mathbb{P}(\theta) + \sum_{r=1}^{R} \log \mathbb{P}(\boldsymbol{n}_r|N_r, \boldsymbol{\alpha}(r)') - \sum_{r=1}^{R} \log \mathbb{P}(\boldsymbol{n}_r|N_r, \boldsymbol{\alpha}(r)),$$

where the sums over all runs $r$ are calculated only for the relevant $r$, i.e. the run $r$ in case of updating $\epsilon_r$, all the runs that correspond to a certain individual $i$ in the case of updating $s_i$, and all the runs that correspond to a certain sequencing type $m$ in the case of updating $\sigma_m$ and $\gamma_{mc}$.

Since $t_c$ are discrete values, the posterior distribution can be evaluated analytically as the integral for normalization turns into a sum. We can therefore sample $t_c$ directly from its posterior distribution using Gibbs sampling:

$$t_c \sim \frac{\mathbb{P}(\rho_c|t_c, v, m_{\mathrm{Y}}, m_X)\mathbb{P}(t_c|\boldsymbol{\pi})}{\sum_{\xi\in\{\mathrm{A,Y,X,D}\}} \mathbb{P}(\rho_c|\xi, v, m_{\mathrm{Y}}, m_X)\mathbb{P}(\xi|\boldsymbol{\pi})},$$

Although $s_i$ are also discrete and could also be updated using Gibbs sampling, we use Metropolis-Hastings instead because Gibbs requires the calculation of the likelihood for all seven possible karyotypes, which is computationally expensive. However, since the aneuploid karyotypes are less likely, we use custom proposal probabilities $q = (0.4, 0.4, 0.04, 0.04, 0.04, 0.04, 0.04)$ for proposing karyotypes $\mathrm{XY, XX, X, XXY, XYY, XXX, XXYY}$, respectively, and correct for this in the Hastings ratio.

### 4 Parameter initialization

To reduce the computational time of the MCMC, we initialize the model parameters to good starting values prior to running the MCMC. We start by initializing $\boldsymbol{s}$ using principal component analysis (PCA) on the normalized depth, similar to the algorithm used by Nursyifa et al. (2022). We create a matrix $\hat{\boldsymbol{\Gamma}}^{(m)}$ for each sequencing type $m$ with elements $[\hat{\boldsymbol{\Gamma}}]_{ic}^{(m)}$ given by

$$[\hat{\boldsymbol{\Gamma}}]_{ic}^{(m)} = \frac{\sum_{r,\mathcal{M}(r)=m} n_{rc}}{\sum_{r,\mathcal{M}(r)=m} L_{rc}} \cdot \frac{\sum_{r,\mathcal{M}(r)=m}\sum_{k=1}^{K} L_{rk}}{\sum_{r,\mathcal{M}(r)=m}\sum_{k=1}^{K} n_{rk}},$$

where $L_{rc}$ is the reference statistics (e.g. length) for sequencing run $r$ of scaffold $c$ and $K$ corresponds to a user-defined number of largest scaffolds that are used to normalize the depth (default 5). The term $\mathcal{M}(r) = m$ indicates if a run $r$ corresponds to sequencing method $m$. For each sequencing method $m$, let us define by $\mathcal{S}_m$ the set of individuals indices that were sequenced with this sequencing type and by $J = |\mathcal{S}_m|$ its length. The dimensions of $\hat{\boldsymbol{\Gamma}}^{(m)}$ then given by $J \times C$.

We then center this matrix by subtracting the mean normalized depth:

$$[\hat{\boldsymbol{\Gamma}}]_{ic}^{(m)} = [\hat{\boldsymbol{\Gamma}}]_{ic}^{(m)} - \frac{1}{J}\sum_{i\in\mathcal{S}_m} [\hat{\boldsymbol{\Gamma}}]_{ic}^{(m)}$$

We then do a k-means clustering with $K = 2$ components for each scaffold $c$ and sequencing type $m$ separately. For each such clustering, we calculate the area of overlap of two normal distributions characterized by the mean and variance of the two inferred clusters. If the area of overlap is smaller than $\exp(-5)$, we consider the two clusters as significantly different. If more than two scaffolds are significant, we denote by $\hat{\boldsymbol{\Gamma}}^{(m)*}$ the matrix containing only the significant scaffolds and do a PCA on $\hat{\boldsymbol{\Gamma}}^{(m)*}$. We then use the two first principal components as input for another k-means clustering with $K = 2$ components (assuming only karyotypes XY and XX). For each $m = 1, \ldots, M$ sequencing types, we use separate cluster means but shared cluster assignments per individual. If no scaffold has a significant area of overlap, we simply use the scaffold with the smallest area of overlap. In this case, or if a single scaffold is significant, we don't do PCA but simply use the clusters inferred by the k-means clustering on that scaffold.

We thus get for each individual an assignment to which cluster it belongs, however, we do not yet know which cluster corresponds to which sex karyotype.

Let us define by $\hat{\pi}_{imc}$ the normalized counts:

$$\hat{\pi}_{imc} = \frac{\sum_{r,\mathcal{I}(r)=i,\mathcal{M}(r)=m} n_{rc}}{2\sum_{r,\mathcal{I}(r)=i,\mathcal{M}(r)=m} N_r},$$

where $\mathcal{I}(r) = i$ indicates if a run $r$ corresponds to individual $i$, and $\mathcal{M}(r) = m$ indicates if a run $r$ corresponds to sequencing method $m$. Here, we assume that most scaffolds are autosomes with a ploidy of two. We start by assigning all individuals in the first cluster to XY and all individuals in the second cluster to XX. For each scaffold $c$ and sequencing method $m$, we set $\hat{\rho}_c = \mathrm{R}$ where $\mathrm{R} \in \{\frac{1}{2}, 0.1, 0.9\}$ and evaluate the linear model

$$\hat{\boldsymbol{\pi}}_{mc} = \boldsymbol{p}_c \hat{\gamma}_{mc} + \epsilon_{mc},$$

where $\hat{\boldsymbol{\pi}}_{mc} = (\hat{\pi}_{\mathcal{S}_1 mc}, \ldots, \hat{\pi}_{\mathcal{S}_J mc})$ is a vector of normalized counts for each individual in set $\mathcal{S}_m$, $\boldsymbol{p}_c = (p_{\mathcal{S}_1 c}, \ldots, p_{\mathcal{S}_J c})$ are the ploidies that are given by the assignment of $s$ and $\rho_c$ for the same individuals, and $\epsilon_{mc}$ is an error term following $\epsilon_{mc} \sim \mathcal{N}(0, \sigma^2_{mc})$. Then

$$\hat{\boldsymbol{\pi}}_{mc} \sim \mathcal{N}(\boldsymbol{p}_c \hat{\gamma}_{mc}, \sigma^2_{mc}),$$

such that we can solve for $\hat{\gamma}_{mc}$ by

$$\hat{\gamma}_{mc} = \frac{1}{J} \sum_{i \in \mathcal{S}_m} \frac{\hat{\pi}_{imc}}{p_{ic}}.$$

The sum of squares is given by

$$S(\hat{\gamma}_{mc}) = \sum_{i \in \mathcal{S}_m} (\hat{\pi}_{imc} - \hat{\gamma}_{mc} p_{ic})^2.$$

We calculate $S(\hat{\gamma}_{mc})$ for every value of $R$, and set $\hat{\rho}_c$ to the $R$ minimizing the sum of squares over all sequencing types $\sum_{m=1}^{M} S(\hat{\gamma}_{mc})$.

We then swap the assignment of individuals by assigning all individuals in the first cluster to XX and all individuals in the second cluster to XY. We repeat the linear regressions as described above. Finally, we choose the configuration that minimizes the overall sum of squares $\sum_{c=1}^{C} \sum_{m=1}^{M} S(\hat{\gamma}_{mc})$, and set all $\hat{\gamma}_{mc}$, $\hat{s}_i$ and $\hat{\rho}_c$ accordingly, where we normalize $\sum_{c=1}^{C} \hat{\gamma}_{mc} = 1$.

The remaining parameters $(\epsilon_1, \ldots, \epsilon_R)$ and $(\sigma_1, \ldots, \sigma_M)$ are set to a small values, $1^{-5}$ and $\exp(-10)$, respectively.

#### 4.1 Extended initialization

We noticed that we could break the algorithm when deliberately initializing all $\boldsymbol{\rho} = \frac{1}{2}$ (autosomes) and $\boldsymbol{s}$ to random values. The algorithm would then sometimes find a solution where the assignment of $\boldsymbol{s}$ is switched, i.e. all males become females and vice versa, and compensate this with an assignment of $\boldsymbol{t}$ and $\boldsymbol{\gamma}$ such that a stable, but sub-optimal solution resulted. To get from this solution to the correct solution would require crossing a region of very low likelihoods, which is very unlikely to happen. As a solution, we implemented the following "extended" initialization procedure that takes place during the first burnins:

1. To simplify the model, we start by allowing only euploid karyotypes XY and XX and $\rho_c = \{\frac{1}{2}, 0.1, 0.9\}$. We run $K$ iterations and then write the current state, i.e. the values of all parameters as well as their proposal width, to a file. In addition, we store the current likelihood.

2. We then switch all karyotypes: For all $i$, if the average $\bar{s}_i = \frac{1}{K}\sum_{k=1}^{K} s_i^{(k)}$ so far was $\bar{s}_i < \frac{1}{2}$, set $s_i = 1$, else set $s_i = 0$. We run another $K$ iterations where we fix $\boldsymbol{s}$ at these values such that the other parameters can adjust to this configuration.

3. Then we run another $K$ iterations where we also update $\boldsymbol{s}$.

4. We evaluate which configuration was better: If the likelihood of the old configuration (prior to switching) is higher than the current likelihood, we read the state file and re-set all parameters and proposal widths to the corresponding values. Else we continue with the current parameter values and proposal widths. We now allow all values for $\rho_c$ while still restricting $\boldsymbol{s}$ to the euploid karyotypes and run $K$ iterations.

5. Finally, we also allow aneuploid karyotypes. We run another $J$ iterations of burnin.

We use $K = 2000$ by default, which results in $4K = 8000$ iterations of extended initialization.

### 5 Supplementary Figures

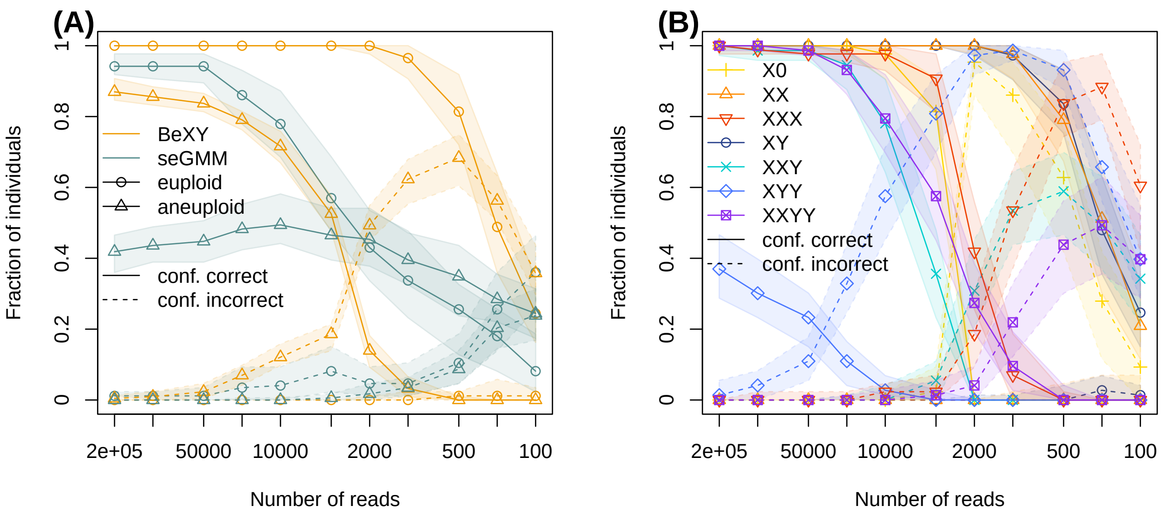


Figure 2: Power to sex individuals. Shown are the median (line with symbols) and 98% confidence interval (shaded area) of the fraction of individuals classified as confidently correct (solid line) and confidently incorrect (dashed line) across 100 downsampling replicates. (A) The power of `BeXY` compared to `seGMM` to classify both euploid and aneuploid karyotypes using a stringent prior of $a = \frac{1}{440}$. (B) The same results of `BeXY` as in A, but plotted by karyotype.

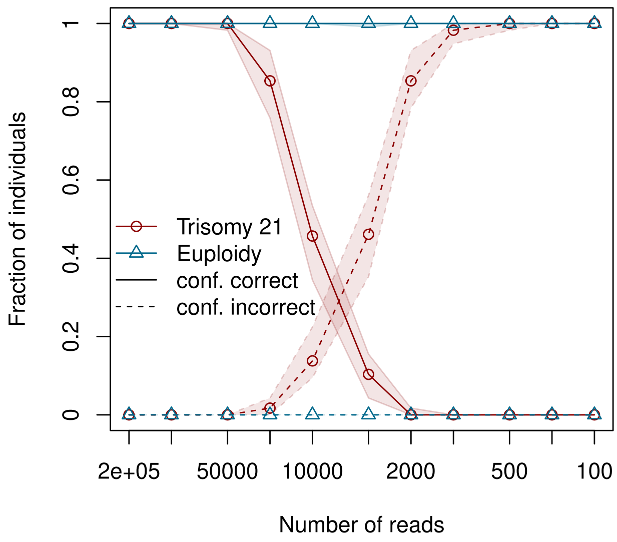


Figure 3: Power to infer autosomal trisomies. The median (line with symbols) and 98% confidence interval (shaded area) of the fraction of individuals classified as confidently correct (solid line) and confidently incorrect (dashed line) as a function of sequencing depth across 100 downsampling replicates when using the prior $\pi = \frac{1}{1000}$. The red line corresponds to the classification of individuals with simulated trisomy 21, while the blue line corresponds to the classification of euploid individuals.

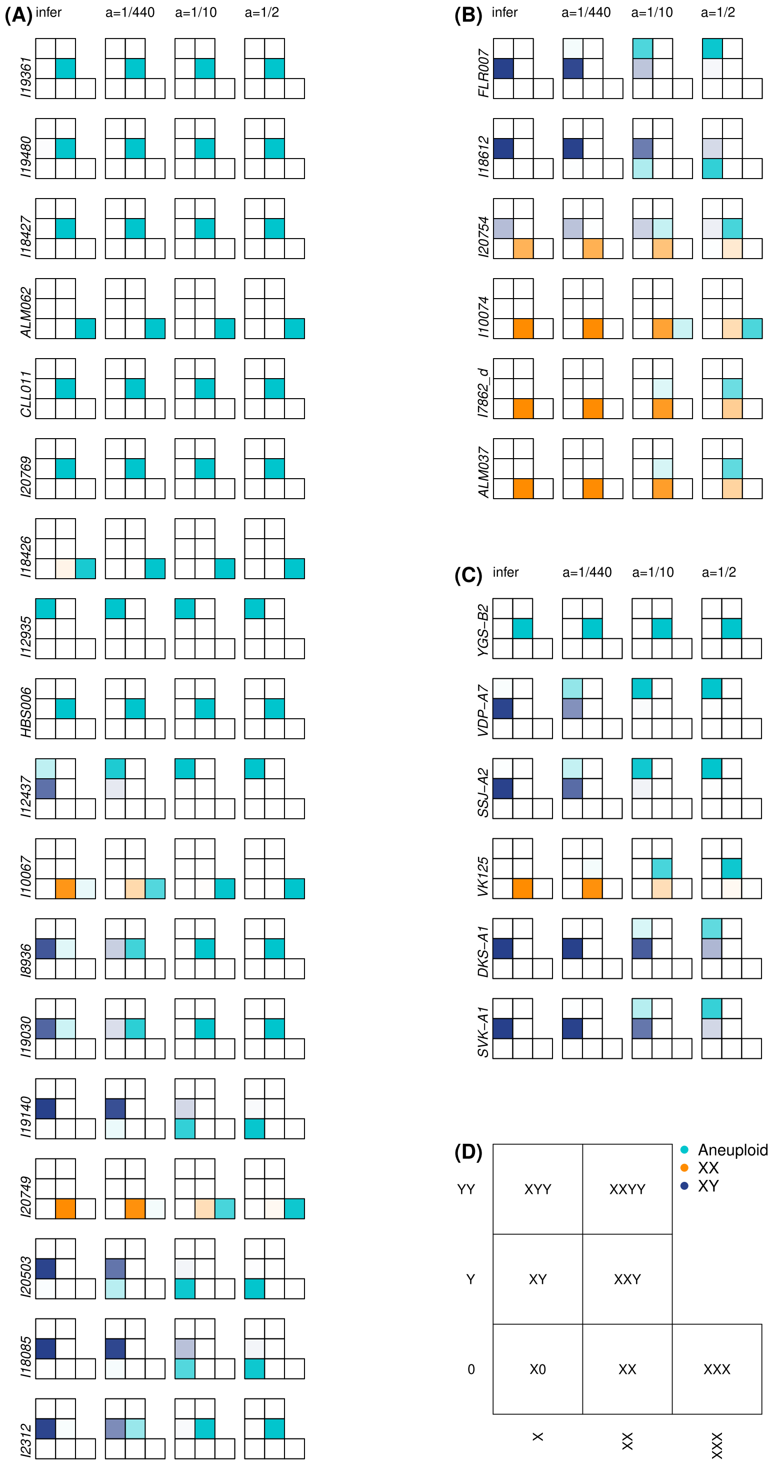


Figure 4: The effect of the prior choice $a$ on aneuploid sex karyotypes. The first column of panels A, B and C corresponds to the results of the task `infer`, which is shown in Figure 5 of the main text. The other three columns correspond to the results from task `sex`, where the parameter $a$ was set to $\frac{1}{440}$, $\frac{1}{10}$ and $\frac{1}{2}$, respectively. (A,B) The posterior probabilities of sex karyotypes of all 1240k target enrichment capture samples classified as aneuploid in at least one case, split into two parts for a better layout. The samples ALM062.A0101.TF1.1, CLL011.A0101.TF1.1, ALM037.A0101.TF1.1 and I2312_all_d were abbreviated as ALM062, CLL011, ALM037 and I2312, respectively. (C) The posterior probabilities of sex karyotypes for all samples of the ancient WGS data set classified as aneuploid in at least one case. (D) A legend on how to read a 7-cell matrix. The x-axis corresponds to having one, two or three copies of the X-chromosome and the y-axis corresponds to having zero, one or two copies of the Y-chromosome. The shading intensity represents the posterior probability of that particular sex karyotype.

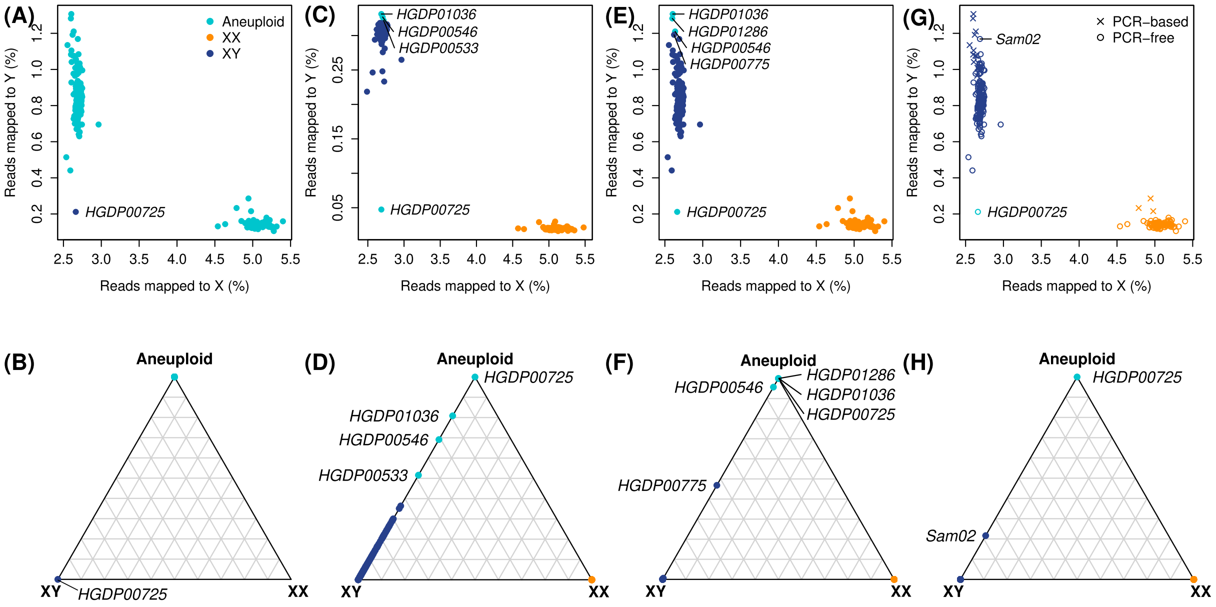


Figure 5: Sex karyotype classification of modern human samples. The top row shows the percentage of reads mapped to Y against the percentage of reads mapped to X. The individuals are colored according to the posterior mode of the sex karyotype. The second shows the posterior probabilities for each sample to have an XY, XX or aneuploid sex karyotype as a ternary plot. (A, B) `BeXY` ran with the task `sex` on the original CRAM-files, using parameters $\boldsymbol{\rho}$, $\gamma_m$ and $\sigma_m$ inferred on the full set of 954 ancient WGS samples. (C, D) `BeXY` ran with the task `sex` on the CRAM-files filtered for a minimum mapping quality of 30, using parameters $\boldsymbol{\rho}$, $\gamma_m$ and $\sigma_m$ inferred on the full set of 954 ancient WGS samples. (E, F) `BeXY` ran with the task `infer` on the original CRAM-files. (G, H) `BeXY` ran with the task `infer` on the original CRAM-files, but treating PCR-based and PCR-free as two different sequencing types. The plotting symbol corresponds to the library preparation protocol.
